# Fully Human Bifunctional Intrabodies Achieve Graded Reduction of Intracellular Tau and Rescue Survival of *MAPT* Mutation iPSC-derived Neurons

**DOI:** 10.1101/2024.05.28.596248

**Authors:** Lianna D’Brant, Natasha Rugenstein, Su Kyoung Na, Michael J. Miller, Timothy F Czajka, Nicole Trudeau, Emily Fitz, Lindsay Tomaszek, Elizabeth S. Fisher, Ethan Mash, Shona Joy, Steven Lotz, Susan Borden, Katherine Stevens, Susan K. Goderie, Yue Wang, Taylor Bertucci, Celeste M. Karch, Sally Temple, David C. Butler

**Affiliations:** Neural Stem Cell Institute, One Discovery Drive, Rensselaer NY 12144; Department of Psychiatry and Knight Alzheimer Disease Research Center, Washington University in St Louis, St Louis, MO, USA

## Abstract

Tau protein aggregation is a hallmark of several neurodegenerative diseases, including Alzheimer’s disease, frontotemporal dementia (FTD) and progressive supranuclear palsy (PSP), spurring development of tau-lowering therapeutic strategies. Here, we report fully human bifunctional anti-tau-PEST intrabodies that bind the mid-domain of tau to block aggregation and degrade tau via the proteasome using the ornithine decarboxylase (ODC) PEST degron. They effectively reduced tau protein in human iPSC-derived cortical neurons in 2D cultures and 3D organoids, including those with the disease-associated tau mutations R5L, N279K, R406W, and V337M. Anti-tau-hPEST intrabodies facilitated efficient ubiquitin-independent proteolysis, in contrast to tau-lowering approaches that rely on the cell’s ubiquitination system. Importantly, they counteracted the proteasome impairment observed in V337M patient-derived cortical neurons and significantly improved neuronal survival. By serial mutagenesis, we created variants of the PEST degron that achieved graded levels of tau reduction. Moderate reduction was as effective as high reduction against tau V337M-induced neural cell death.

## Introduction

Tauopathies encompass a range of neurodegenerative dementia-causing diseases characterized by the accumulation of insoluble tau protein in neurons and glia. There are over twenty clinicopathological diseases associated with abnormal tau, including Alzheimer’s disease (AD), progressive supranuclear palsy (PSP) and familial forms such as frontotemporal lobar degeneration with tau inclusions (FTLD-tau) and due to mutations in the *MAPT* gene that encodes tau.^1^ Under normal conditions, tau protein is abundant within neurons and is low in astrocytes and oligodendrocytes, but with disease tau can accumulate in glial cells, contributing to pathology spread ^2–5^ In AD and other tauopathies, tau protein appears to lose its ability to bind to microtubules and becomes mis-localized to the somatodendritic compartment of neurons.^6^ During this process, tau may become abnormally hyperphosphorylated and forms insoluble aggregates, such as straight filaments and paired helical filaments (PHF) which comprise neurofibrillary tangles (NFTs).^7^ The presence of ubiquitin-positive tau in neurofibrillary tangles (NFTs) correlates with the severity of AD-related dementia.^8, 9^ Despite the presence of ubiquitin on the molecule, which normally spurs proteasomal degradation, this abnormal tau is poorly degraded.^8^

Considerable evidence indicates that lowering tau protein can be neuroprotective in animal models and in induced pluripotent stem cell (iPSC)-derived neuronal models.^10–15^ Hence, significant effort has been expended on developing tau-lowering strategies. The majority of these have focused on lowering extracellular tau, with the concept that by disrupting neuron-neuron spread, pathology spread can be curtailed. However, to date, therapeutics aimed at lowering extracellular tau have not proven effective. The approaches used, including passive immunotherapy, active immunization, anti-aggregation agents, and modulation of tau phosphorylation, have yet to demonstrate efficacy in altering disease course.^16–20^ One explanation is that the vast majority of pathological tau protein is building up inside neural cells, and that targeting extracellular tau does not address this source of cellular toxicity. Intrabodies, or intracellular antibodies, represent a promising avenue for targeting tau inside cells. Our approach involves combining a tau-targeting antibody with a PEST degron, a polypeptide sequence enriched in proline (P), glutamic acid (E), serine (S), and threonine (T), to achieve effective intracellular tau lowering via proteasomal degradation. This method can be used to degrade both normal and misfolded tau.^21, 22^

Intrabodies are small antibody fragments derived from the variable region (Fv) of full-length immunoglobulin.^23^ They retain many advantages of conventional antibodies, including high specificity and affinity for target epitopes, but lack the Fc domain that is responsible for mediating inflammatory reactions.^23^ Intrabodies are powerful tools to target putative pathogenic proteins because they can be selected, engineered, and delivered as genes by strategies such as AAV delivery. Bifunctional intrabodies that include a degron offer a promising approach for selectively targeting and lowering specific proteins by harnessing the cell’s protein degradation machinery. Our previous studies have demonstrated the efficacy of bifunctional intrabodies in reducing aggregation and stimulating the degradation of abnormal proteins associated with Huntington’s disease and Parkinson’s disease.^23–28^ In this study, we describe the development and characterization of fully human bifunctional intrabodies engineered to target and degrade intracellular tau.

In the normal adult brain, six tau isoforms are expressed, and there are approximately equal amounts of tau protein with three or four repeat microtubule binding domains (3R or 4R tau). ^29, 30^ Depending on tauopathy, the abnormal tau accumulations may consist of predominantly 3R or 4R tau or a mixture. Moreover, cryo-EM has revealed that the underlying structure of abnormal tau aggregation is distinct in different tauopathies and influenced by post-translational modifications.^31–34^ To limit tau aggregation, we aim to reduce the level of tau protein. We created bifunctional molecules by fusing an anti-tau antibody binding domain to the ornithine decarboxylase (ODC) PEST degron that promotes degradation through the proteasome via ubiquitin-independent proteolysis.^35, 36^ The PEST degron binds to the lid (19S) of the 26S proteasome^36, 37^ and provides a flexible tether allowing the intrabody and its bound protein to enter the proteasome for degradation.^36^ Recognizing the potential drawbacks of excessive tau reduction, we have serially mutated the human PEST degron and identified sequences that alter the degradation efficiency. This approach, we term ’programmable target antigen proteolysis’ (PTAP), enables precise modulation of tau protein levels. In this study, we demonstrate that anti-tau-PEST intrabodies effectively reduced both overexpressed and endogenous tau levels in neuronal cell lines and iPSC-derived cortical organoids and significantly increased neuronal survival. Moreover, they improved proteasome function in iPSC-derived *MAPT* mutation neurons, maintaining efficacy even under conditions of proteasome impairment, a characteristic feature of aging and neurodegenerative diseases such as FTD and AD.^38–40^ Overall, our findings reveal the therapeutic potential of fully human bifunctional intrabodies as a ‘one and done’ gene therapy strategy to achieve targeted tau reduction in tauopathies, offering promise for the development of disease-modifying therapies for these devastating neurodegenerative diseases.

## Results

### Bifunctional anti-tau-PEST intrabodies with mouse ODC PEST effectively lower overexpressed tau in murine immortalized cells and human 2D iPSC-derived cortical neurons

To identify intrabodies that efficiently target tau protein for degradation via the proteasome, we developed a series of human anti-tau-intrabodies using 17 anti-tau single-chain Fvs (scFvs) selected from a large non-immunized phage-display library of 6.7 x 10^9^ members.^41^ The library was originally modified to enhance the selection of soluble intracellular scFvs specifically targeting tau amino acids (aa) 151-441.^42^ This central region of the tau protein, which contains the proline rich regions and microtubule binding domains, plays an important role in tau aggregation and pathology.^20^ Hence, this is an attractive region to target with an intrabody in order to block tau aggregation.

Given that this phage display library has produced several scFvs with nanomolar to picomolar affinity,^41–43^ we hypothesized that the intrabodies we generated would exhibit sufficient intracellular affinity to direct tau to the proteasome for degradation when fused to the mouse ODC PEST degron (mPEST) (**Figures 1A, B).** The 17 anti-tau-mPEST intrabodies were co-transfected into the ST14A rat striatal progenitor cell line^44^ with a GFP-tagged 4R tau construct (eGFP-Tau- (0N4R))^6^ (**Figure 1C)**. This initial screen revealed 11 candidate anti-tau-mPEST intrabodies that effectively lowered the steady-state eGFP-Tau-(0N4R) levels relative to empty-vector control (EV CON) **(Supplemental Figure S1)**. Anti-tau intrabodies V-mPEST, N-mPEST, and F-mPEST significantly (** p<0.01) reduced eGFP-Tau-(0N4R) by ∼78%, 70%, and 74% respectively compared to EV CON (**Figures 1D-F).** The anti-tau intrabodies were initially shown to bind tau epitopes 151-274, 368-391, and 402-411.^42^ To confirm this for each of our lead candidates, we performed epitope mapping. We generated a series of GFP-tau deletion constructs ranging from tau amino acids 151-274 to 151-441, enabling us to use GFP as the primary readout for target engagement. The anti-tau-mPEST intrabodies and GFP-Tau deletion constructs were co-transfected into ST14A cells and harvested at 48 hours for western blotting. We found that F- mPEST bound to an epitope within tau amino acids 368-391 **(Supplemental Figure S1E).** V- mPEST and N-mPEST did bind to any of the deletion constructs, suggesting that they may detect a conformational epitope within tau.

**Figure 1.**
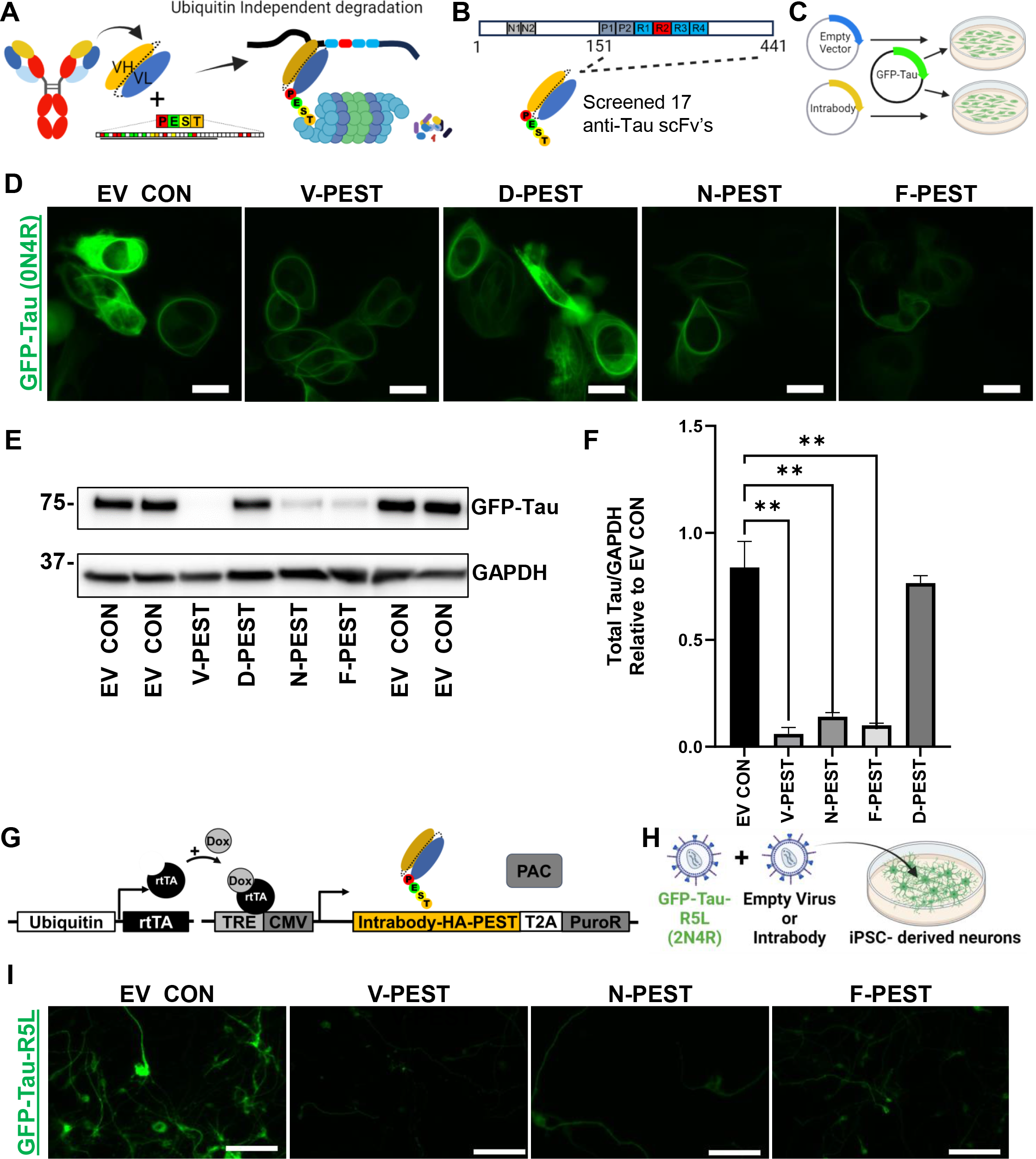
**Bifunctional anti-Tau intrabodies significantly lower GFP-Tau A)** Schematic of intrabody isolation and engineering. Recombinant intracellular antibodies, known as Intrabodies, contain the variable heavy (VH) and variable light (VL) antigen binding domains of conventional full-length antibodies. Fusion of the ODC mPEST degron facilitates ubiquitin-independent degradation of the intrabody and its bound antigen. **B)** Tau intrabodies were selected against amino acids 151-441, which contains the proline-rich regions and microtubule-binding domains. **C)** Experimental design for intrabody selection. Murine ST14A cells were co-transfected with GFP-Tau and anti-tau-mPEST intrabodies or EV CON. **D)** Fluorescence images of GFP-Tau illustrate a reduction in GFP-Tau signal in V, N and F intrabody-treated cells. Scale bar = 20 µm **E)** Corresponding immunoblot data of GFP-Tau protein, quantified by densitometric analysis in **F);** Data represent mean ± SEM. n=3; One-way ANOVA was performed followed by Tukey’s multiple comparisons post-hoc tests (**p<0.01). **G)** Schematic of the dox-inducible intrabody lentiviral construct. **H)** Experimental design for intrabody screening in iPSC-derived neurons. **I)** Fluorescence images illustrating GFP-Tau signal reduction in intrabody-treated cells. Scale bar = 20 µm.

We next investigated the impact of anti-tau intrabodies on tau levels in human iPSC-derived cortical neurons. To assess this, and to determine whether intrabodies could lower mutated tau that causes FTD-tau, we generated an overexpression plasmid containing full length (2N4R) GFP-tagged tau with the R5L mutation as the target by site-directed mutagenesis of GFP-Tau 2N4R tau. The R5L mutation is associated with PSP and forms aggregated, insoluble tau protein predominantly consisting of four-repeat isoforms.^45^ This mutation modifies the N terminal structure of tau and its ability to condense on the surface of microtubules.^46^ Overexpression of GFP-Tau-R5L in ST14A cells revealed the presence of extensive puncta, membrane blebbing, and cytotoxicity not seen with the GFP-Tau control **(Supplemental Figure S3A).** We utilized a dox-inducible lentiviral system in which iPSC-derived 2D cortical neurons were co-transduced with reverse tetracycline transactivator (rtTA) regulating expression of a bifunctional HA-tagged anti-tau-mPEST intrabody (either V-mPEST, N-mPEST, F-mPEST) or an empty virus control (EV CON) together with GFP-tau-R5L (2N4R) (**Figure 1G, 1H).** Subsequently, we induced the intrabody expression by treating the cultures with 2 µg/mL of doxycycline (DOX). Following 72 hours of intrabody expression, we observed a significant reduction in GFP-tau-R5L expression with all three anti-Tau-mPEST intrabody treatments (**Figure 1I).**

### Bifunctional anti-tau-mPEST intrabodies lower endogenous tau in 3D iPSC-derived cortical organoids

Having shown that the anti-tau-mPEST intrabodies can effectively lower overexpressed tau in 2D cultures of enriched human cortical neurons, we next evaluated whether the intrabodies could lower overexpressed tau in the context of organoids as 3D models of cellularly complex cerebral cortical tissue. To do this we used cerebral cortical patterned organoids generated from iPSCs using our established methods (**Supplemental Figure 2**).^47, 48^ By 4 months, the resulting organoids include the major cortical deep and upper layer neurons, subsets of inhibitory neurons, astrocytes, and oligodendrocyte progenitor cells.

At 42 days of differentiation, we co-transduced the organoids with dox-inducible V-mPEST intrabodies or EV CON, reverse tetracycline transactivator (rtTA) and GFP-Tau-R5L (2N4R) **(Supplemental Figure S3B, S3C).** Following 21 days of treatment with DOX (2 µg/mL) to induce intrabody expression, the organoids were live imaged by time-lapse confocal microscopy for GFP-Tau-R5L expression (note that on intrabody-treated organoids, the level of GFP was significantly lowered, hence the exposure time had to be increased 10-fold for visualization) **(Supplemental Figure S3D)**. In the EV CON-treated organoids, cytotoxicity was observed, demonstrated by swelling in the axonal processes followed by blebbing and cell death **(Supplemental Figure S3E, Supplemental Video 1)**. The blebbing was similar to that we observed in ST14A cultured cells. In contrast, the cells in V-mPEST intrabody treated organoids were dynamic and healthy without obvious blebbing or death **(Supplemental Figure S3E, Supplemental Video 2)**.

These studies demonstrated that the bifunctional intrabodies could bind and lower overexpressed GFP-labeled tau. To establish whether the intrabodies could efficiently and safely reduce endogenous tau, 4-month organoids without tau mutations were co-transduced with lentiviral vectors expressing rtTA and an inducible anti-tau intrabody (V-mPEST, N-mPEST, or F- mPEST) or EV CON (**Figure 2A).** Following 21 days of treatment with 2 µg/mL of DOX, V-mPEST, N-mPEST, and F-mPEST significantly (* p<0.05, ** p<0.01) were expressed and reduced endogenous total tau protein by 55.3%, 57.9%, and 38.82%, respectively, compared to EV CON (**Figures 2B,C).** We next evaluated lowering endogenous tau using organoids derived from iPSCs from patients with heterozygous missense *MAPT* (CGG to TGG) mutation in exon thirteen, which encodes R406W, and their CRISPR corrected controls. Individuals with the R406W mutation exhibit a disease course similar to Alzheimer’s disease (AD), characterized by initial memory loss, a late onset, prolonged disease duration and the accumulation of AD-like tau PHFs consisting of the 6 brain isoforms.^49–53^ 4-month-old organoids with R406W or CRISPR corrected isogenic controls were co-transduced with reverse tetracycline transactivator (rtTA) and either anti-tau intrabodies or EV CON (**Figure 2D).** Following 60 days of treatment with 2 µg/mL of DOX, the resulting 6-month organoids were cryostat sectioned and immunostained for tau. V-mPEST, N-mPEST, and F-mPEST significantly reduced endogenous total tau quantified by immunofluorescence compared to EV CON in both WT and R406W organoids (**Figure 2E, F),** confirmed by western blot analysis (**Figure 2G, H).** In contrast, the number of MAP2ab+ neurons appeared to be consistent across treatment groups.

**Figure 2.**
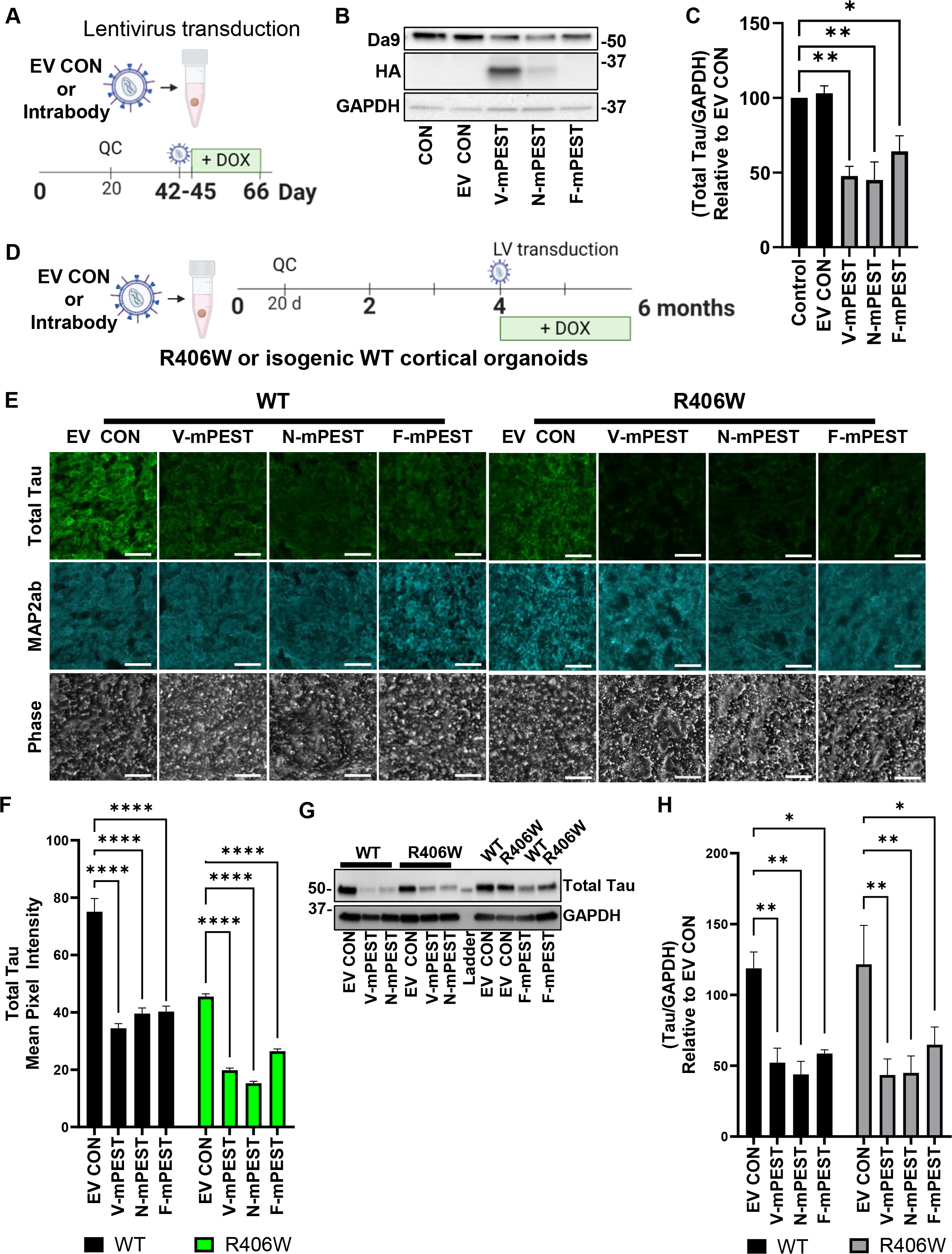
**Bifunctional anti-tau intrabodies reduce endogenous tau in human cortical organoids A)** Experimental design for the treatment of iPSC-derived organoids (healthy donors). **B)** Representative western blot of total tau (DA9) and intrabody (HA) levels following 21 days of treatment; GAPDH as loading control quantified by densitometry in **C);** bars represent total tau mean optical density values relative to GAPDH loading control. The data are presented relative to EV CON. Mean ± SEM; One-way ANOVA with Tukey’s post-hoc test (* p<0.05, **p< 0.01). **D)** Experimental design: Four-month-old cortical organoids derived from isogenic R406W and CRISPR-corrected controls were transduced with either anti-tau intrabodies (V-mPEST, N- mPEST, F-mPEST), or EV CON. At six months, organoids were either fixed and cryosectioned for immunofluorescent staining with antibodies to total tau and MAP2ab or harvested for western blotting. **E)** Representative images of immunostained cryostat sections. Scale bar = 20 μm. **F)** Total tau mean pixel intensity. **G)** Representative western blots quantified by densitometry in **H)**; Data represents mean ± SEM, n =3; Two-way ANOVA with Dunnett’s multiple comparisons post- hoc test. * p<0.05, ** p<0.01, **** p<0.0001.

### Development of human tau-lowering intrabodies

In pursuit of our primary goal to create a safe and effective reagent for human use, we conducted a comparative analysis between the mouse and the human ODC PEST (hPEST) degron, a region rich in proline (P), glutamic acid (E), aspartic acid (D), serine (S), and threonine (T) (**Figure 3A)**. There is 75% homology between the mPEST and hPEST degron. We test its functionality, we substituted the human (hPEST) for the mouse degron in the V-intrabody constructs, then compared their ability to lower intracellular tau. We co-transfected HEK293 cells with GFP-Tau (0N4R) and either V-mPEST, V-hPEST, or EV CON. After 48 hours of treatment, cells were live imaged for GFP expression (**Figure 3B)** and then lysed for biochemical analysis. Both V-hPEST and V-mPEST significantly lowered the mean GFP pixel intensity compared to EV CON (**** p < 0.0001) (**Figure 3C)** confirmed by western blot analysis of total tau protein (** p<0.01) (**Figure 3D)**. Importantly, there was no significant difference between V-hPEST and V-mPEST.

**Figure 3.**
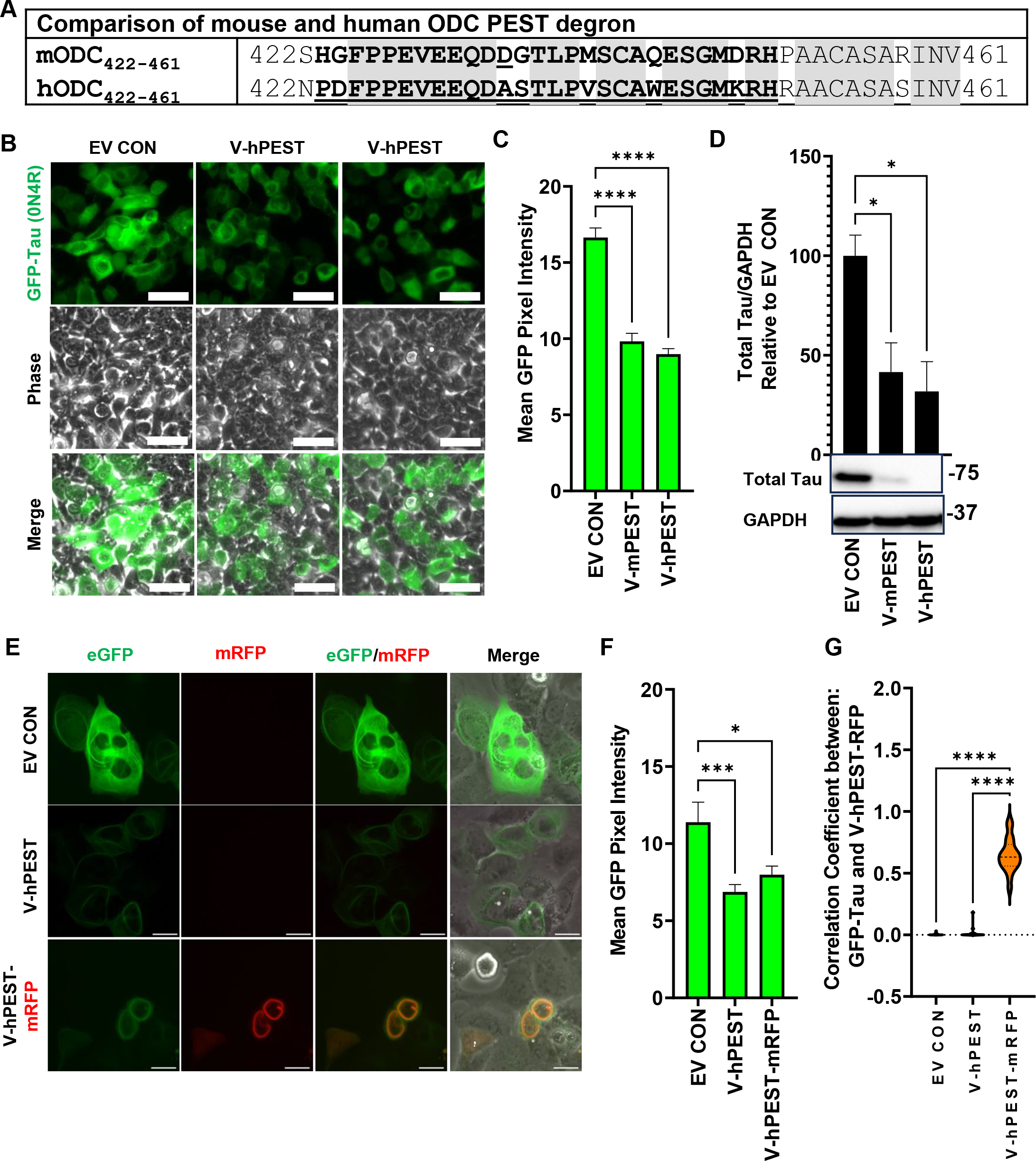
**Bifunctional anti-tau intrabodies with a human PEST degron exhibit comparable tau reduction to those with a mouse PEST degron. A)** Sequence comparison of the human and mouse ODC PEST degron; the PEST region is indicated in bold. Conserved areas are shaded grey; non-conserved regions have no shading. The C-terminal 40 amino acids of the mouse ODC degron shares 75% (30/40) homology with human ODC; the consensus PEST sequence is underlined. **B)** HEK293 cells were co-transfected with GFP-Tau (0N4R) and empty vector control (EV CON) or anti-tau intrabodies with either mouse or human ODC PEST degrons. Representative live cell images show GFP-Tau reduction in V-mPEST and V-hPEST treated cells at 72 hours compared to EV CON. Scale bar = 20 μm. **C)** Quantification of GFP-Tau, mean GFP pixel intensity. **D)** Quantitative western blotting of total tau (DA9) normalized to the GAPDH protein loading control. The data are presented relative to EV CON; One-way ANOVA with Tukey’s post-hoc test; (* p < 0.05). **E)** ST14A cells were co-transfected with GFP-Tau (0N4R) and either EV CON, V-hPEST, or V-hPEST-mRFP. Representative live cell images show GFP-Tau reduction in V-hPEST and V-hPEST-RFP treated cells at 72 hours compared to EV CON. Scale bar = 20 μm. **F)** Quantification of GFP-Tau; mean GFP pixel intensity. **G)** Pearson correlation coefficient for colocalization of GFP (tau) and mRFP (intrabody).

We next wanted to assess the localization and functionality of an intrabody targeting tau in live cells. To assess intrabody localization with tau, we developed a live cell anti-tau-intrabody- reporter by fusing mRFP, along with a flexible Gly4Ser4 linker, to the C-terminus of the intrabody. The Gly4Ser4 linker allows the intrabody and mRFP to fold independently. This approach enables us to track the intrabody’s distribution and confirm its interaction with tau in real time. Furthermore, by comparing the performance of V-hPEST with and without the mRFP tag, we sought to determine whether the fluorescent reporter affects the intrabody’s ability to reduce tau levels. Here we co-transfected GFP-Tau with either EV CON, V-hPEST, or V-hPEST∼mRFP in rat ST14A cells. We observed a significant reduction in GFP expression with both V-hPEST (*** p < 0.001) and V-hPEST∼mRFP (* p< 0.05) (**Figure 3E,F)**. There was no significant difference between V- hPEST and V-hPEST∼mRFP, which suggests that the mRFP tag does not interfere with the intrabody’s ability to degrade tau. Using Image J, we observed a significant Pearson’s correlation coefficient (0.60, **** p<0.0001) between GFP-Tau and V-hPESTmRFP, indicating a high degree of co-localization between the intrabody and tau in the cells (**Figure 3G).**

### Targeted degradation of tau in human astrocytes with anti-tau intrabodies

While *MAPT* gene expression is normally low in macroglial cells, abnormal tau protein frequently accumulates in tauopathies, impairing glial cell function and contributing to the spread of pathology. Hence, targeting the tau protein to reduce accumulated tau is likely to be more effective than targeting the *MAPT* RNA in glial cells. To confirm the functionality of anti-tau-hPEST intrabodies in this setting, we co-transduced human iPSC-derived astrocytes with an inducible lentiviral vector expressing GFP-Tau-V33M-P2A-mRFP1, along with either V-hPEST, N-hPEST, or an empty vector control (EV-CON). The P2A sequence allows GFP-Tau-V337M and mRFP1 to be produced from the same mRNA and cleaved via a P2A peptide during translation. Doxycycline was added to the culture medium three times per week at 2 µg/mL to induce the transgene expression. After 14 days of treatment, live-cell imaging was performed to assess GFP-Tau- V337M expression **(Supplemental Figure 4A).** Both V-hPEST and N-hPEST significantly (p < 0.01) reduced GFP-Tau-V337M compared to the EV CON **(Supplemental Figure 4B).** These findings demonstrate that bifunctional anti-tau-hPEST intrabodies effectively degrade mutant tau within astrocytes.

### Bifunctional anti-tau-PEST intrabodies lower tau via ubiquitin-independent proteolysis

Protein degradation is regulated through the Ubiquitin-Proteasome System (UPS) which conjugates ubiquitin molecules onto proteins that are destined to be degraded. Ubiquitination is a multistep process involving ubiquitin-activating (E1), ubiquitin-conjugating (E2), and ubiquitin-ligating (E3) enzymes (**Figure 4A).** To confirm that bifunctional anti-tau-PEST intrabodies facilitate ubiquitin-independent rather than ubiquitin-dependent proteolysis, we used the small molecule PYR-41 to inhibit E1 ligation and epoxomicin (Epox) to inhibit the 26S proteasome (**Figure 4A).**

**Figure 4.**
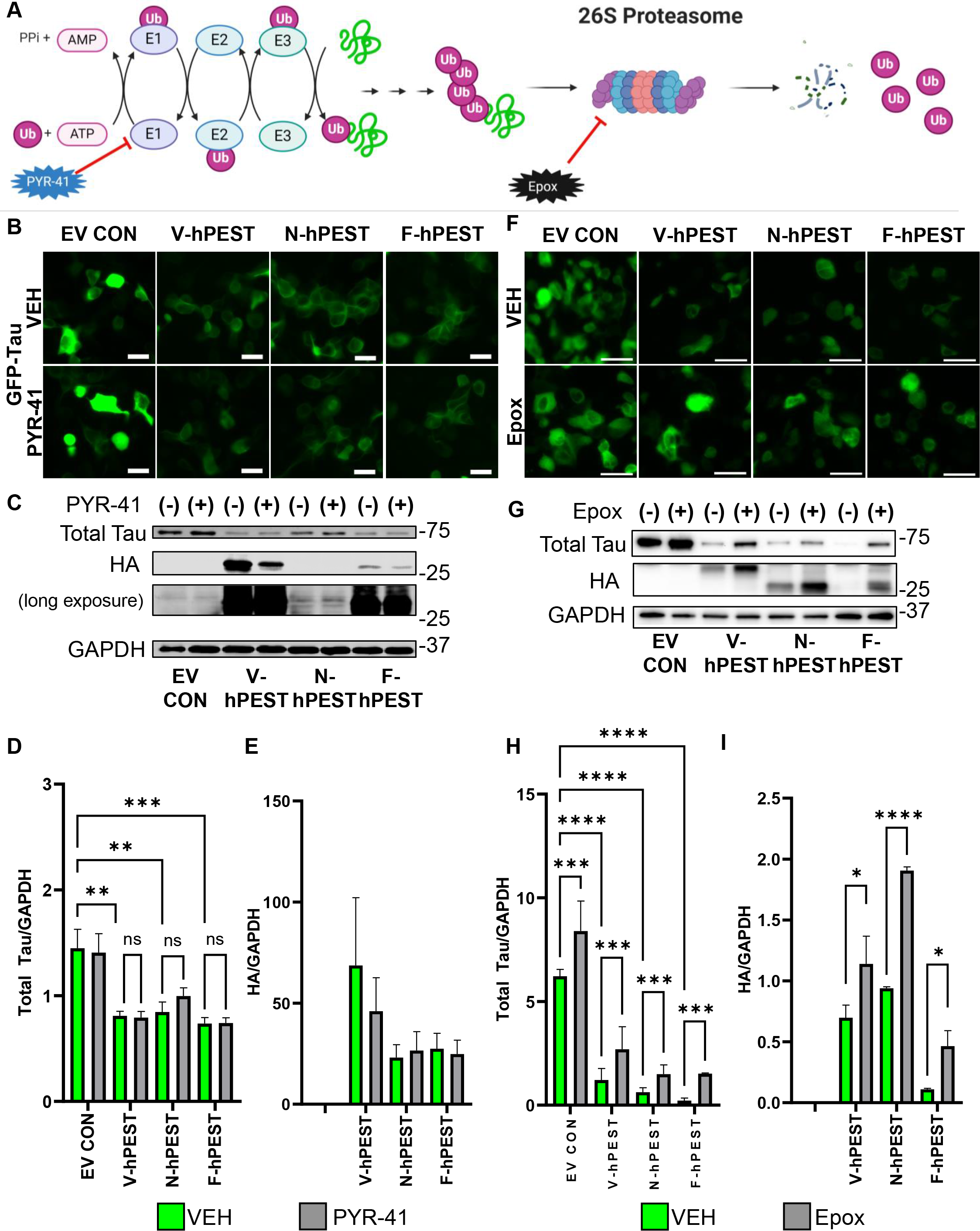
**Bifunctional anti-tau Intrabodies lower Tau via the Proteasome. A)** Schematic of the ubiquitin-proteasome system and drug inhibition targets. HEK293 cells were co-transfected with GFP-Tau (0N4R) and anti-tau intrabodies (V-hPEST, N- hPEST, F-hPEST), or EV CON, and treated with PYR-41 to block E1 or Epoxomicin to inhibit 26S proteasome degradation. **B-E)** <u>PYR-41 inhibition of ubiquitin-activating enzyme, E1</u> (50 µM, 4 hours): **B)** Representative live cell images. **C)** Representative western blots quantified by densitometry in **D);** total tau (DA9) normalized to GAPDH protein loading control. Total tau is significantly lower in intrabody treated groups compared to EV CON and is not impacted by PYR- 41 treatment, and in **E)** intrabody levels (HA) normalized to GAPDH protein loading control; PYR- 41 treatment does not alter intrabody protein levels. **F-I)** <u>Proteasome inhibition:</u> Proteasome inhibition with Epoxomicin (Epox; 10µM, 16 hours). **F)** Representative live images show an increase in GFP-Tau expression in Epox treated cells compared to VEH (DMSO) control. **G)** Representative western blots show tau and intrabody (HA) protein levels are increased in HA treated groups compared to VEH control, quantified by densitometry in **H);** total tau normalized to GAPDH as protein loading control and in **I);** intrabody levels (HA) normalized to GAPDH as protein loading control. Two-way ANOVA with Tukey’s post-hoc test; ( * p<0.05, **p< 0.01, ***p<0.001, **** p<0.0001.)

We first tested our hypothesis that the bifunctional intrabodies operated via ubiquitin-independent proteolysis. HEK293 cells were co-transfected with GFP-Tau (0N4R) and either V- hPEST, N-hPEST, F-hPEST, or an empty vector control (EV CON) construct. Post-transfection, cells were treated with PYR-41 (50 µM, 4 hours), an E1 ligase inhibitor, or dimethyl sulfoxide (DMSO) as a vehicle control (VEH). If tau degradation was ubiquitin-dependent, GFP-Tau levels would increase following PYR-41 treatment. However, as expected, V-hPEST, N-hPEST, and F- hPEST significantly lowered GFP-Tau compared to EV CON (**Figure 4B-D).** Importantly, tau levels and intrabody (HA) levels did not significantly differ between intrabody treatment groups (VEH and PYR-41). Due to the short half-life of intrabodies, longer exposure times were required to effectively detect HA-tagged N-hPEST intrabody levels (**Figure 4C)**, quantified in **Figure 4E**.

Our next goal was to confirm that the tau-lowering effects of anti-tau-hPEST intrabodies are mediated by proteasome-dependent proteolysis. We co-transfected HEK293 cells with GFP-Tau and V-hPEST, N-hPEST, F-hPEST, or empty vector control (EV CON) constructs. Post-transfection, cells were treated with Epox (10 µM for 16 hours), a selective 26S proteasome inhibitor, or vehicle control (DMSO). As expected, V-hPEST, N-hPEST, and F-hPEST significantly reduced GFP-Tau protein levels compared to the EV CON group (p < 0.0001) but reduction was blocked by Epox treatment, as observed through live cell imaging and western blot analysis (p < 0.001) compared to the VEH control (**Figure 4F-H)**. Intrabody levels, assessed using HA tags were minimally detected in VEH-treated cells due to their short intracellular half-life, while proteasome inhibition with Epox significantly increased intrabody levels relative to the VEH control group (**Figure 4I).**

### Pharmacokinetics and half-life of intrabody and tau protein in cellular models

Unmodified single-chain variable fragment (scFv) intrabodies have been reported to have half-lives ranging from 8 to 20 hours depending on several factors including their specific sequence, stability, and the cellular environment in which they are expressed.^54–56^ Based on previous studies with GFP-mPEST,^57, 58^ we hypothesized that the half-life of anti-tau-hPEST intrabodies would be approximately 2 hours. To determine intrabody half-life, we co-transfected ST14A cells with GFP-Tau (0N4R) and either V-hPEST, V-hPEST-Scrambled (V-hSCR; inactive degron control), or EV CON. 48 hours after transfection, cells were treated with CHX (100 μg/mL), a global protein translation inhibitor,^59^ for 0, 2, 3, or 6 hours (**Figure 5A).** The half-life was calculated according to the formula: 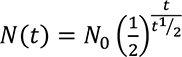 where *N*(*t*) = quantity of the substance remaining, and *N*_0_ = the initial quantity of the substance and *t* = time elapsed. The half-life of GFP-Tau in EV CON treated cells was 4.89 hours (**Figure 5B,C)**. *In vitro*, tau has been reported to have a half-life of between 5-60 hours using immortalized cells,^60^ which is considerably shorter than the half-life of tau in post-mitotic iPSC-derived cortical neurons, which has been measured at ∼6.74 days.^18^ V-hPEST shortened the half-life of GFP-Tau to approximately 2.93 hours compared to EV CON (**Figure 5B,C)**. In contrast, V-hSCR, which contains an inactive PEST degron, did not shorten the half-life of GFP-Tau (**Figure 5B,C)**. V-hPEST and V-hPEST-Scrambled intrabody half-lives were 1.75 and 3 hours, respectively (**Figure 5D).**

**Figure 5.**
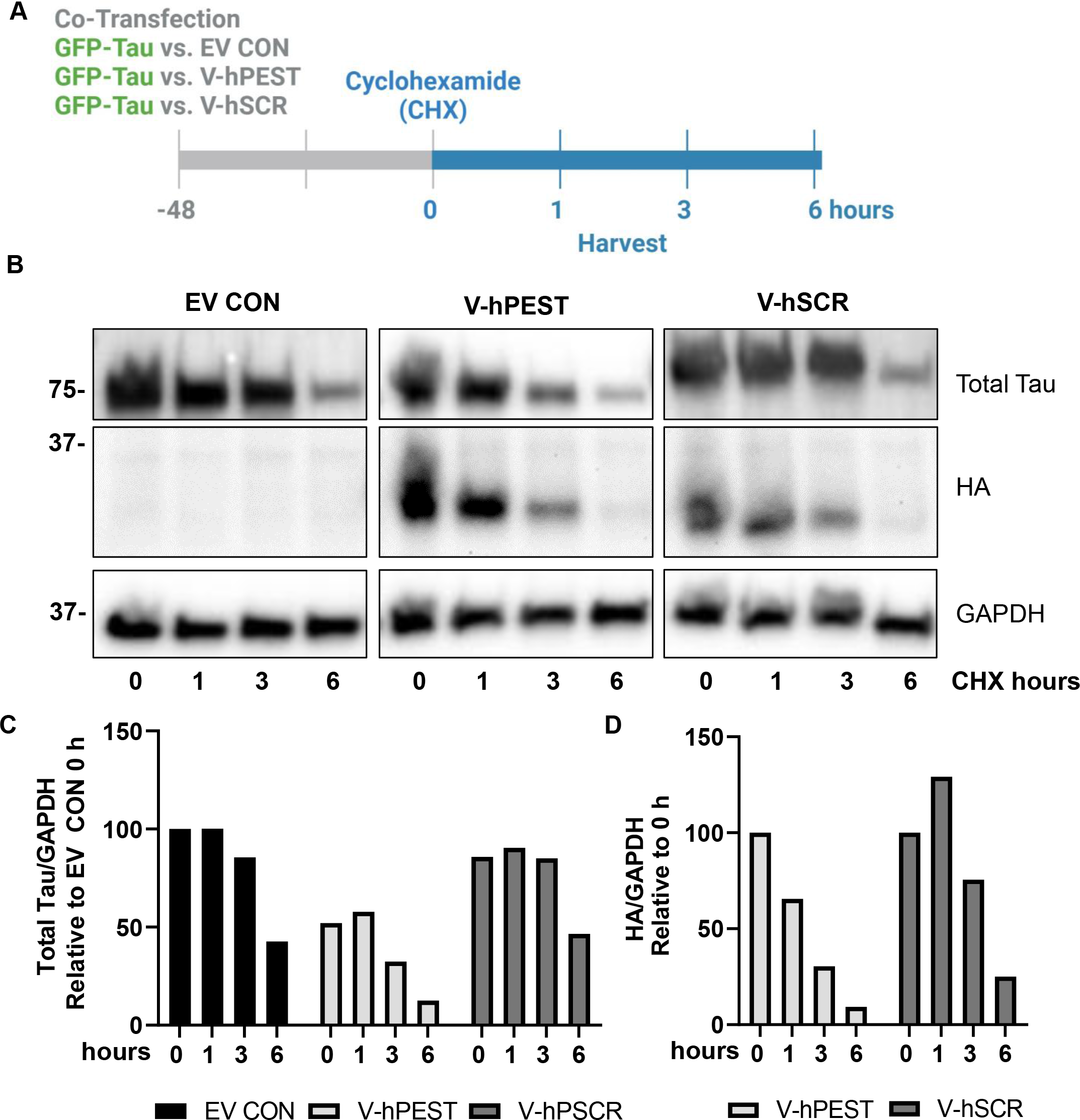
**The half-life of tau is reduced by V-hPEST compared to EV CON. A)** Experimental design. Murine immortalized ST14A neural precursor cells were co-transfected with GFP-Tau (0N4R) and either V-hPEST, V-hPEST-Scrambled (V-hSCR; inactive degron control), or EV CON. 48 hours after transfection, transfected cells were treated with cycloheximide (CHX; 100μg/mL), a global protein translation inhibitor, for 0, 1, 3, or 6 hours. **B)** Representative western blot of total tau protein (DA9) and intrabody quantified by densitometry in **C)**; total tau normalized to EV CON at time point zero and in **D);** intrabody levels (HA) normalized to time point zero.

### Bifunctional anti-tau-PEST intrabodies counteract proteasome impairment in patient-derived cortical neurons

One concern regarding utilizing the proteasome to reduce tau protein is the reported impaired proteasome function in brain regions affected by tauopathy, for example in AD.^61, 62^ To determine whether proteasome impairment impacts the efficacy of anti-tau intrabodies, we first showed that proteasome function was impaired in human iPSC-derived neurons with the *MAPT* V337M mutation, which is associated with behavioral variant FTD.^63^ Patients with the V337M mutation develop NFTs of 3R and 4R tau which are similar to those found in AD.^63^ The lines used in this analysis were obtained from two donors carrying the V337M *MAPT* mutation and their corresponding CRISPR-corrected controls.^64^ iPSC-derived cortical neurons with the *MAPT* V337M mutation and CRISPR-corrected control neurons at 90 days were co-transduced with lentiviral vectors expressing a fluorescently tagged ubiquitin reporter (Ub^G76V^-eGFP) to visualize ubiquitin-dependent proteolysis, serving as a test for proteasome function in living cells, along with either anti-tau-PEST intrabodies (V-hPEST, N-hPEST, F-hPEST) or an empty vector control (EV CON) (**Figure 6A,B)**.^65–67^ Under normal conditions, Ub^G76V^-GFP is rapidly degraded by the 26S proteasome but, conversely, it builds up in cells when the proteasome activity is impaired. Our data indicate a significant impairment in proteasome function in *MAPT* V337M mutant cells compared to isogenic controls (p < 0.0001) (**Figure 6C,D).** However, we found that bifunctional anti-tau-hPEST intrabody expression can significantly (p < 0.0001) counteract proteasome impairment due to the *MAPT* V337M mutation (**Figure 6C,D**).

**Figure 6.**
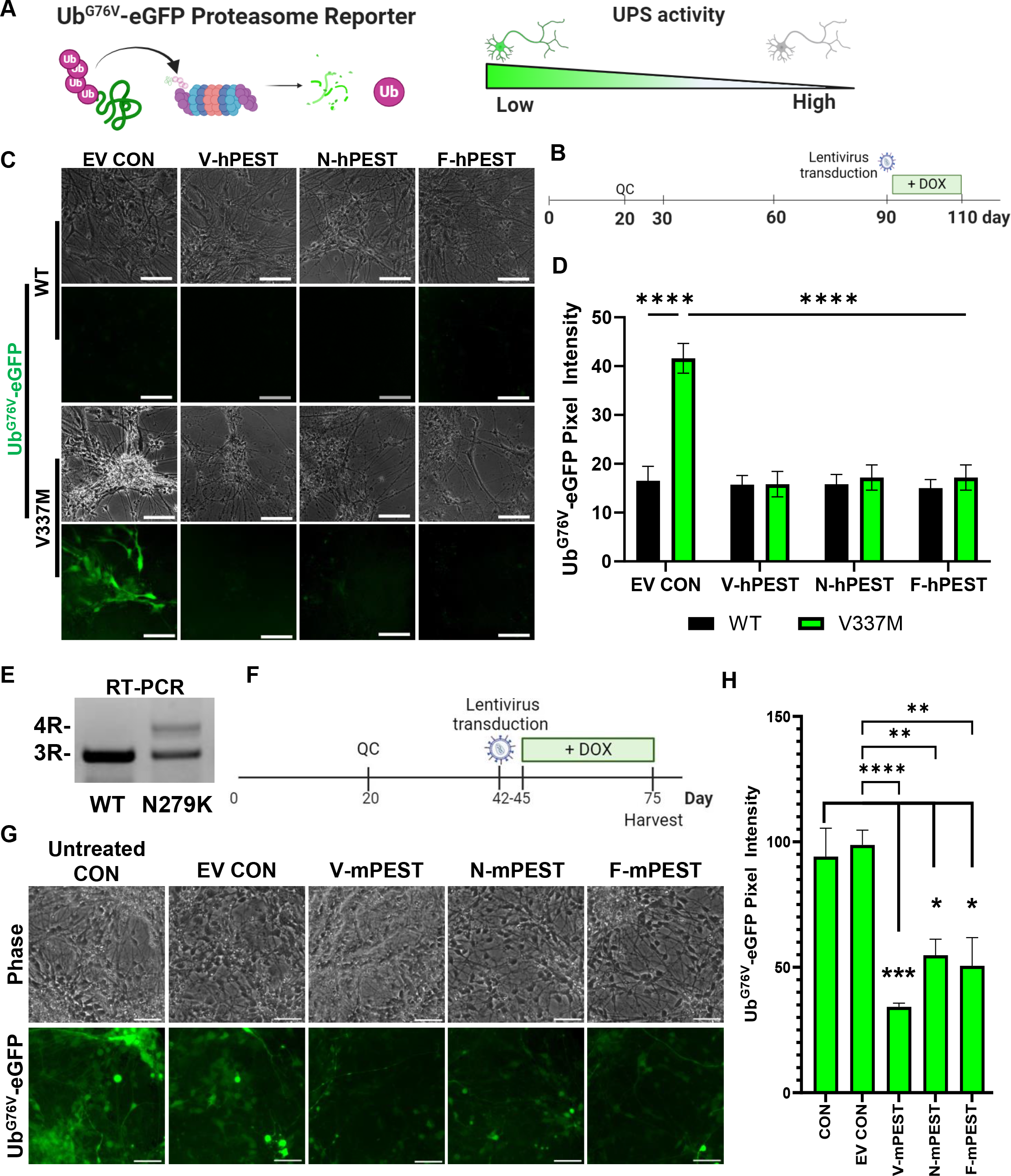
**Anti-tau intrabodies counteract proteasome impairment in iPSC derived cortical neurons. A)** Schematic of Ub^G76V^-eGFP reporter. **B)** Representative images of Isogenic V337M and CRISPR corrected controls transduced with lentiviral vectors expressing Ub^G76V^-eGFP and either EV CON, V-hPEST, N-hPEST, or F-hPEST according to the experimental design **C).** Representative live cell images show Ub^G76V^-eGFP in V337M *MAPT* mutation cortical neurons. Scale bar = 50 µm. **D)** Quantification of Ub^G76V^-eGFP reporter mean eGFP pixel intensity; mean ± SEM. Two-way ANOVA with Tukey’s post-hoc test (****p<0.0001). **E)** N279K 2D cortical cultures mature to express a 1:1 ratio of 3R:4R tau. **F)** Experimental design. **G)** Representative live cell images. Scale bar = 50 µm. **H)** Quantification of Ub^G76V^-eGFP reporter in N279K cortical cultures following 30 days of treatment; mean ± SEM. One-way ANOVA with Tukey’s post-hoc test. (*p< 0.05, **p<0.01, ***p<0.001, **** p<0.0001.)

Because iPSC-derived cortical neurons fail to achieve the mature level of 4R tau production even after extensive culture periods,^64^ we next generated iPSCs with the *MAPT* mutation N279K by CRISPR engineering **(Supplemental Figure 5).** This mutation significantly increases expression of 4R tau in patients and leads to FTD with Parkinsonism.^68^ By qPCR, we confirmed a near 1:1 ratio of 3R and 4R tau in homozygous N279K neurons after 60 days *in vitro* (**Figure 6E).** These neurons were co-transduced with Ub^G76V^-eGFP and either anti-tau-mPEST intrabodies (V-mPEST, N-mPEST, F-mPEST) or an empty vector control (EV CON) (**Figure 6F).**^65–67^ Similar to our observations in *MAPT* V337M treated cortical cultures, delivery of anti-tau- mPEST intrabodies significantly reduced Ub^G76V^-eGFP expression in *MAPT* N279K cortical neuron cultures (**Figure 6G,H)**, indicating their ability to reduce proteasome impairment in the context of both 3R and 4R tau.

### Partial reduction of tau with anti-tau PTAP intrabodies effectively improves survival of MAPT mutation neurons

Given tau’s multiple cellular functions, complete tau reduction could have undesirable consequences. To regulate the level of tau reduction achieved with anti-tau intrabodies, PTAP variants (V1 and V9) were generated by serially mutating critical residues within the human ODC PEST degron (Figure 7A**, Supplemental Figure 6A,B**). The impact of these mutations was then tested by assessing GFP-Tau degradation efficiency in ST14A cells **(Supplemental Figure 6C)**. Select constructs were then delivered to iPSC-derived cortical *MAPT* V337M mutation before the onset of mutation-induced cell death, which occurs at around 90 days (**Figure 7B).** Anti-tau- hPTAP intrabody administration resulted in different levels of tau reduction, assessed by immunofluorescent staining (**Figure 7C,D)**. Excitingly, anti-tau-hPTAP intrabodies, F-hPEST-V1, F-hPEST-V9, and N-hPEST-V9 all significantly improved the viability of *MAPT* V337M treated cortical cultures similar to corrected control (V337V) levels (**Figure 7E).** The protection achieved with anti-tau-hPTAP F-hPEST-V1 (∼50% in V337M neurons) was similar to the protection achieved with the F-hPEST-degron high reducer (70% in V337M neurons) (**Figure 7C-E).** This indicates that even partial reduction of tau can mitigate the toxic effects of tau accumulation.

**Figure 7.**
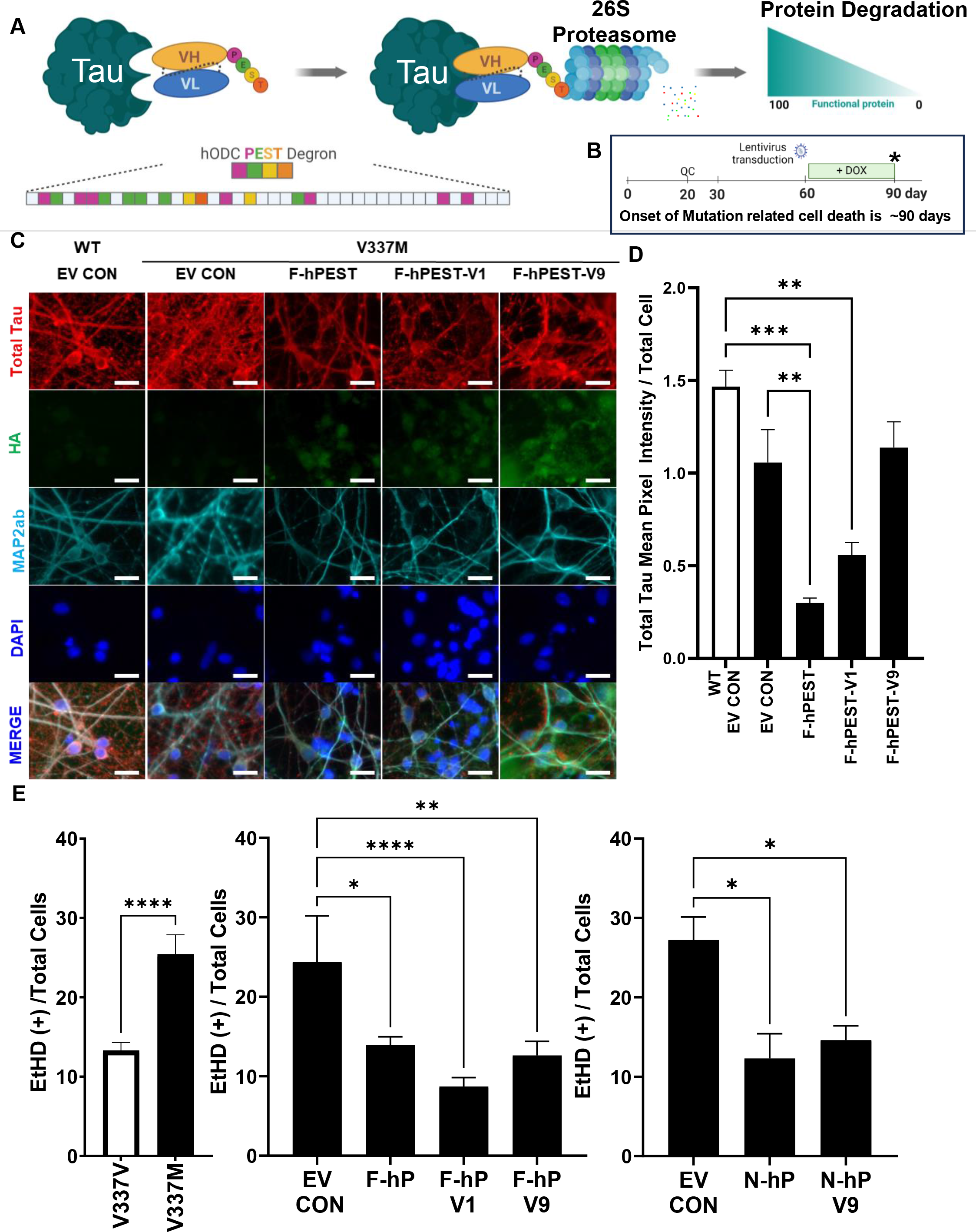
Anti-tau intrabodies with PTAP variants providing high (hPEST) or moderate (V9) tau reduction lower cell death to isogenic WT control levels. A) Schematic of PTAP technology. Modifications to the hODC PEST degron result in differential levels of protein degradation. B) Experimental design. 60-day-old V337M cortical cultures were transduced. At 90- days cultures were assessed for viability or fixed for immunofluorescence staining immunostaining with antibodies to: total tau, intrabody (HA), or MAP2ab. C) Representative images of immunostaining. Scale bar = 20 µm. D) Quantification of total tau mean pixel intensity relative to total cells. E) At 90 days, cell death was detected by ethidium homodimer staining and quantified with CellProfiler^TM^. Mean ± SEM. One-way ANOVA with Tukey’s post-hoc test (*p< 0.05, **p<0.01, ***p<0.001, **** p<0.0001).

## Discussion

Tauopathies represent a group of neurodegenerative disorders characterized by the abnormal accumulation of tau protein in neurons and glia, leading to various forms of dementia. Despite extensive research, therapeutic interventions targeting intracellular tau accumulation have remained elusive. In response to this pressing challenge, our study introduces a novel therapeutic approach: bifunctional anti-tau proteasomal targeting intrabodies designed to bind and reduce tau levels within the cytoplasm. Envisioned as a gene therapy, this innovative strategy offers the potential for a stable, long-term solution to mitigate intracellular tau pathology. By harnessing the cell’s protein degradation machinery, these intrabodies hold promise as a ’one and done’ gene therapy, capable of preventing tau accumulation and aggregation in neurons and glia over the long-term. The approach is distinct from protein lowering approaches that reduce tau via ubiquitin-dependent pathways, or ASO approaches that require repeated administration. By reducing tau toxicity and enhancing neural cell proteostasis and health, tau lowering via bifunctional intrabodies holds significant potential for clinical translation, offering hope for alleviating the burden of dementia in tauopathy patients.

### Efficacy and Mechanism of Action

Recent advancements in intrabody engineering have vastly improved the versatility and efficacy of intrabodies beyond the neutralization of intracellular gene products. Misfolded proteins are generally degraded through two major pathways: the ubiquitin-proteasome system (UPS) and the autophagy-lysosomal proteolysis pathway.^69^ Several pioneering studies illustrated that intracellular proteins can be retargeted to various cellular compartments using intrabodies.^70–72^ Based on these studies, bifunctional intrabodies were designed to selectively direct their target proteins to the UPS.^73, 74^ Melchionna and Cattaneo demonstrated that tau could be targeted to the proteasome for degradation with an anti-tau scFv intrabody fused to IκBα in N2A immortalized cells and SHSY-5Y neuroblastoma cells. ^73^ Their anti-tau intrabody reduced the levels of truncated tau, (amino acids 151-422 of full-length 2N4R tau), and endogenous full-length tau by 70-90%. However, this is not an optimal therapeutic strategy due to potential side effects caused by the requirement for a biologically active substrate, TNFα, to initiate protein degradation.

Our group developed bifunctional intrabodies that can direct misfolded proteins to the proteasome without the need for additional substrates, thus avoiding undesirable off-target effects.^26, 28, 75, 76^ Fusion of the PEST degron from mouse ornithine decarboxylase (amino acids 422-461) to an anti-huntingtin-scFv-C4 or to an anti-synuclein-VH14 resulted in protein reduction of huntingtin or α-synuclein by 80-90%^26^ and 60%,^25^ respectively. Here, through a series of *in vitro* experiments utilizing neural cell models, we developed and characterized anti-tau-PEST intrabodies that are capable of targeting tau to the proteasome for clearance in both murine and human model systems. Our experiments, coupled with live cell imaging and colocalization analysis, confirm the specificity of intracellular tau binding by the intrabodies. Moreover, we discovered that despite several differences in amino acid sequence, the mPEST and hPEST degrons behave similarly in anti-tau bifunctional intrabodies. We found that the intrabodies can degrade 3R and 4R tau with the mPEST degron in human iPSC-derived organoids and 2D cultures (**Figures 1-3** **and Supplemental Figures 1-3).** Conversely, with the hPEST degron, the intrabodies were able to effectively degrade tau in murine cells (**Figure 3E, F)**. This cross-species utility aids translational studies. Importantly, our findings suggest that these intrabodies exert their effects via the proteasome, as evidenced by the reversal of tau reduction upon treatment with a proteasome inhibitor. We have observed similar results with our bifunctional anti-synuclein intrabody VH14 using the hPEST degron and the mPEST degron.^75, 76^ In line with our *in vitro* findings, a recent *in vivo* study utilized a *Drosophila* split-luciferase-based sensor of tau-tau interaction, termed tau^LUM^, to quantify tau multimerization. Transgenic expression of our anti-tau intrabody significantly reduced tau multimerization and cell death in tau^LUM^ flies.^77^ Hence, these bifunctional intrabodies not only facilitate tau proteasomal degradation but also prevent the formation of tau aggregates, likely because they target the central domain of tau that is important for aggregation, highlighting their multi-faceted therapeutic potential.

### Benefit of Ubiquitin-Independent Proteasomal Degradation

The ubiquitin-dependent proteasome system can be leveraged to target proteins for clearance. In humans, there are only two ubiquitin-activating enzymes (E1s), ∼40 ubiquitin-conjugating enzymes (E2s), and over 600 ubiquitin-ligase enzymes (E3s) that regulate ubiquitination.^78, 79^ During ubiquitin-dependent proteolysis, the C-terminal Gly76 of ubiquitin is reversibly ligated to target proteins or other ubiquitin molecules.^78^ Typically, ubiquitin forms polyubiquitin chains through seven internal lysine (K) residues: K6, K11, K27, K29, K33, K48, and K63. Among these, K48-linked ubiquitination is the most common and primarily triggers proteasomal degradation. Gallardo *et al*., designed a chimeric anti-tau intrabody fused to ubiquitin harboring a lysine K63 mutation to arginine (R) to favor polyubiquitination at lysine K48, which results in substrate degradation by the 26S proteasome.^80^ AAV9 delivery of their HJ8.8-K48R chimeric intrabody, which was performed before overt tauopathy had developed, decreased pathological tau in P301S transgenic mice.^80^ Our approach differs because our bifunctional intrabody degradation is ubiquitin-independent, *i.e.*, it does not rely on the ubiquitination process. This is supported by data showing that inhibiting E1 ubiquitin activation with PYR-41 did not result in increased intrabody and tau protein levels, as seen following proteasome inhibition with epoxomicin. This is significant because the UPS can be perturbed with neurodegenerative disease.^78, 81, 82^ Moreover, approaches that utilize chimeric intrabodies with ubiquitin fusions can inadvertently promote protein aggregation as these fusions can become incorporated into existing protein aggregates. Myeku *et al*., observed the co-localization of Ub^G76V^-eGFP and tau in inclusion bodies within hindbrain neurons in JNPL3:Ub^G76V^-GFP transgenic mice. ^62^

### Bifunctional anti-tau-mPEST and -hPEST intrabodies Alleviate Proteasome Impairment

Protein homeostasis (proteostasis) is the balance between protein synthesis, folding, and degradation.^83^ Impaired proteostasis is hypothesized to contribute to the progression of neurodegeneration in AD and FTLD-tau. The UPS is responsible for the degradation of most proteins, whereas lysosomal proteolysis degrades insoluble aggregated proteins such as tau. Aggregated, misfolded proteins can compromise the function of the 26S proteasome complex, contributing to neurodegeneration.^84^ Corroborating this, impaired proteasome function is reported in affected but not unaffected brain regions in Alzheimer’s disease.^61^ 45% of FTLD-tau patients have abnormal tau inclusion bodies consisting of ubiquitinated and hyperphosphorylated tau, consistent with impaired proteostasis.^85^ Attenuated proteolysis of tau is observed in animal and induced pluripotent stem cell (iPSC) models of tauopathy,^86, 87^ including cells transfected with *MAPT* R406W or V337M tau^88^ and A152T iPSC-derived cortical neurons.^86^ Using the Ub^G76V^- eGFP proteasome reporter, we observed proteasome impairment in 2D *MAPT* N279K and V337M cortical cultures. Notably, our lead bifunctional anti-tau-PEST intrabodies were able to counteract proteasome impairment in N279K and V337M cortical forebrain cultures. We suggest that this is due to the intrabody helping to chaperone and clear misfolded protein, relieving proteostatic stress.

### Tau Lowering reduces Human *MAPT* Mutation Associated Neuronal Death

We established a rigorous cell death assay using neuronal cells derived from patient iPSC lines carrying the disease-causing V337M *MAPT* mutation.^64^ Previous studies have shown A152T *MAPT* mutation neurons are more susceptible to stressors such as rotenone, demonstrating elevated cell death.^86^ This vulnerability to stressors only appeared after cultures were approximately 100 days old. Ehrlich *et al*. recently showed that patient-derived iPSC lines with a *MAPT* V337M mutation produce neurons with enhanced tau fragmentation and phosphorylation, decreased neurite extension, and increased susceptibility to oxidative stress.^58^ Unlike prior A152T and V337M findings,^86, 89^ we observed cell death in V337M cortical cultures at 90 days without the need for additional stressors (**Figure 7)**. At 65 days, viability was similar between isogenic and V337M cultures. Surprisingly, we did not observe a significant increase in cell death when challenging the cells with rotenone; this difference may be due to regional vulnerability. The main difference between the studies is that we generated forebrain neurons, whereas Ehrlich et. al.,^89^ produced midbrain neurons. This is relevant because a subset of patients with the V337M mutation have severe frontal lobe atrophy with a high density of NFTs, pretangles, and neuropil threads^63^ while the substantia nigra in the same patients displayed mild atrophy and NFT pathology.^63^ Our results, combined with clinical data, suggest that there is selective vulnerability in forebrain neurons to the V337M mutation alone without additional stressors. Importantly, our bifunctional anti-tau-hPEST intrabodies were able to counteract cell death in these cultures, even when treated after the onset of cell death, suggesting effectiveness against advanced pathology.

### PTAP potential for optimizing protein degradation levels

One of the challenges in targeting tau is the potential for adverse effects resulting from excessive reduction. To address this concern, we developed PTAP variants by serial mutation of the PEST domain that we found achieve different levels of tau reduction when combined with anti-tau intrabodies. This extends prior studies showing mutations in the degron can alter the degradation of GFP and the intrabody half-life.^90^ We have already demonstrated the value of applying this approach to neurodegenerative disease by *in vitro* and *in vivo* studies with our fully human anti-alpha-synuclein intrabody VH14-hPEST.^75^ The current study applies the PTAP technology to tau lowering. Our results indicate that even partial reduction of tau can effectively protect neurons against cell death, highlighting the importance of fine-tuning tau levels for therapeutic benefit.

### Conclusions

A characteristic of tauopathies is the deposition of tau into insoluble fibrillar aggregates. Our study introduces a promising gene therapy approach for tauopathies using bifunctional anti-tau intrabodies that possess anti-aggregation properties and degrade both 3R and 4R tau isoforms, offering a potential long-term solution to mitigate intracellular tau pathology. This strategy leverages the cell’s protein degradation machinery independently of the ubiquitin pathway, which is important given that the UPS system can be impaired in neurodegenerative disease. The efficacy of these intrabodies in reducing tau levels and protecting neurons from cell death, particularly in V337M cortical cultures, underscores their therapeutic potential. Our study demonstrates the broad applicability of anti-tau-PEST intrabodies across different tauopathy disease contexts, including familial forms of FTLD-tau associated with *MAPT* R5L, V337M, R406W and N279K mutations. Further optimization and preclinical studies will be essential to advance this innovative approach towards clinical applications, providing hope for effective treatments for tau-related neurodegenerative disorders.

### Limitations of the study

Further optimization of intrabody variants and preclinical studies will be important to advance these findings towards clinical translation. Exploring the efficacy of intrabodies in animal models of tauopathy and investigating their long-term effects will be crucial for assessing their therapeutic potential.

## Methods

### Intrabody Construct and Plasmid Design

Tau single-chain variable fragments (scFvs) from Visintin *et al*.,^42^ were synthesized, incorporating either a mouse PEST degron, a human PEST degron, or human PEST degron variants. These constructs were then cloned into pcDNA3.1 (+), pAAV-MCS, and pTetO-FUW expression plasmids. To facilitate intrabody detection, a hemagglutinin (HA) epitope tag (amino acid sequence YPYDVPDYA) was fused to the intrabody C-terminus.^25, 76^ To direct the intrabodies and their cargo to the proteasome, a sequence corresponding to amino acids 422–461 from human ODC (GenBank accession number AH002917.2; NPDFPPEVEEQDASTLPVSCAWESGMKRHRAAC ASASINV) or mouse ODC (GenBank accession number EU684749.1; SHGFPPEVEEQ DDGTLPMSCAQESGMDRHPAACASARINV) was added to the C-terminal end of the HA-tag. To modulate the targeted degradation of the intrabody and its bound antigen, single and compound mutations were introduced within the human PEST degron at specific amino acid positions: P426A/P427A (V1), E428A-E430A-E431A (V2), D433A (V9), S435A (V10), P438A (V12), S440A (V13), C441A (V14), E444A (V15), S445A (V16), K448A (V18). These mutations were synthesized commercially (GenScript, USA). The intrabodies were subcloned into pcDNA3.1 (+) and pTetO-FUW using standard restriction enzyme cloning techniques, following the strategy: EcoRI-NheI-intrabody-NotI-HA-PEST degron-stop-codon-HindIII-XbaI. The intrabodies were subcloned into pAAV at the 5’-XbaI and 3’ HindIII restriction sites according to the following strategy: Nhe1-intrabody-NotI-HA-PEST degron-stop-codon-HindIII. The Nhe1/Xba1 site is lost upon ligation. All expression plasmids were validated by Sanger DNA sequencing (Azenta Life Sciences) and prepared using Nucleobind Xtra Midi Endotoxin-free prep kits (Takara Cat # 740420.5), following the manufacturer’s protocol.

### Lentivirus vector production and viral packaging

Anti-tau intrabodies were cloned into an inducible tetracycline on 3rd generation lentiviral vector, pTet-O-Ngn2-puro, (gift from Marius Wernig; Addgene plasmid # 52047).^91^ The NGN2 cassette was replaced with the tau scFvs and their respective mPEST, hPEST, or hPEST degron variants at the 5’ EcoRI and 3’ Xba1 cloning sites. The tetracycline-on system utilized in this study is dependent on the reverse tetracycline-controlled transactivator, FUW-M2rtTA (rtTA) (a gift from Dr. Rudolf Jaenisch, Addgene plasmid # 20342).^92^ Lentiviruses were produced in 293FT cells (ThermoFisher; Cat # R70007). The intrabody-expressing pTet-O-puro or rtTA plasmids (5.6 µg) were co-transfected with the envelope plasmid pVSV-G (7.1 µg) and the packaging plasmid pCMVd8.9 (14.2 µg) into 293FT cells using 122 µg of transfection-grade linear polyethylenimine hydrochloride (PEI MAX; Polyscience Inc; Cat# 24765). The pVSV-G and pCMVd8.9 plasmids.^93^ After 3 hours, the transfection medium was replaced with fresh medium. Viral-containing media samples were collected at 48 and 72 hours, transferred to sterile 50 mL conical tubes, and centrifuged at 350 x g for 10 minutes. The viral supernatant was filtered through a 0.45 µm cellulose acetate low protein binding filter (Corning, Cat # 09-761-35) and stored at 4°C for 2 to 24 hours before further processing. Viral supernatants were then concentrated using Lenti-X Concentrator (Takara Bio; Cat # 631232) according to the manufacturer’s protocol and resuspended in 1X CMF PBS. Lentivirus titers were determined using a qRT-PCR lentivirus titration kit (ABM, Cat # LV900). For viral transduction, lentiviral vectors at a multiplicity of infection (MOI) of 1 were added to the organoids.

### Cell Culture and Transfection

Undifferentiated ST14A cells, a rat neuronal cell line, and Human Embryonic Kidney (HEK) 293 cells were cultured and transfected with intrabody constructs using polyethylenimine (PEI), following previously described protocols with slight modifications.^75, 76^ Briefly, cells in a 6-well plate were co-transfected with 3 µg of plasmid DNA per well, using JetPEI transfection reagent (Polyplus Transfection Inc); co-transfection: a 3:1 ratio of intrabody expression plasmid or empty vector control (EV CON) to prk5-GFP-Tau plasmid per well. The PEI DNA transfection reagent facilitated transient transfection. Cells were imaged and harvested 72 hours post-transfection.

### Generating 2D forebrain cortical neurons and astrocytes from human iPSCs

Neural precursor cells (NPCs) were provided by the NeuraCell core facility and used to generate 2D cortical neurons, following a modified protocol described by Telezhkin *et al*.^94^ Briefly, iPSCs were cultured to 85-90% confluency in mTeSR1 medium (StemCell Technologies #85850) on 6- well tissue culture (TC) plates (Corning Cat # 3506) coated with Cultrex (R&D Systems Cat # 3434-010-02). Colonies were rinsed twice with 1X PBS (Gibco #14200-075) and then treated with neural induction media supplemented with SLI (Advanced DMEM/F12, Gibco Cat # 12634-028; 2mM Glutamax, Gibco Cat # 35050-061; 2% Neurobrew-21 without RA, Cat # Miltenyi 130-097- 263; 1% Anti-Anti, Gibco Cat # 15240-062; 20uM SB, Tocris Cat # 1614; 200nM LDN, Cat # Tocris 6053; 1uM endo-IWR1, Tocris Cat # 3532). Cells were fed daily from day 0 to day 4. On day 5, endo-IWR1 was removed from the media, and cells were fed daily with neural induction media without endo-IWR1 from day 5 to day 10. On day 10, the cultures were treated with 10 uM Y- 27632 (Tocris #1254) for 1 hour before dissociation using StemPro Accutase (Gibco Cat # A110501) for 20 minutes at 37°C and replated at a 1:2 ratio onto 6-well TC plates coated with Cultrex and cultured in media without SB431542 and LDN193189. Cells were fed daily with neural induction media from day 11 to day 17. Subsequently, cells were either frozen in Cryostor CS10 (Stem cell technologies) or passaged at a 1:2 ratio in preparation for terminal differentiation.

### Neuronal differentiation from cryopreserved NPCs

Day 17 NPCs obtained from NeuraCell were thawed into one well of a Cultrex-coated 6-well plate (Corning Cat # 3506). The cells were then cultured in neural induction media (Advanced DMEM/F12 supplemented with 2mM Glutamax, 2% Neurobrew-21, and 1% Anti-Anti) with daily feeding until day 22. Neuronal differentiation was performed using SCM1 and SCM2 media as described in Telezhkin et al.^94^ On day 23, the cells were plated for neuronal differentiation at a density of 320,000 cells per well of a 6-well plate coated with Cultrex, and cultured in SCM1 medium (Advanced DMEM/F12 (1:1) supplemented with 2mM Glutamax, 2% Neurobrew-21, 1% Antibiotic-Antimycotic, 10 µM DAPT, 10 µM Forskolin, 300 µM GABA, 3 µM CHIR99021, 2 µM PD0332991, 1.8 mM CaCl2, 200 µM ascorbic acid, 1 µM LM22A4 and 10 ng/ml BDNF). The medium was changed by 50% every 2-3 days. On day 31, the medium was exchanged for SCM2 medium (Advanced DMEM/F12 (1:1) and Neurobasal A (Gibco) (50:50) supplemented with 2mM Glutamax, 2% Neurobrew-21, 1% Antibiotic-Antimycotic, 1.8 mM CaCl2, 3 µM CHIR99021, 2 µM

PD 0332991, 200 µM ascorbic acid, 1 µM LM22A4 and 10 ng/ml BDNF), with a 50% medium change every 2-3 days until day 45. Subsequently, there was a full medium change to BrainPhys (StemCell Technologies Cat # 5790) supplemented with 2% Neurobrew-21, 1% Antibiotic-Antimycotic, 10 ng/mL BDNF, and 10ng/mL GDNF (Fuji Cat # 100-02). Cells were fed every 2-3 days with a 50% medium change until being harvested for immunostaining or immunoblotting at 60, 90, and 110 days.

### Astrocyte differentiation

Human astrocytes (ScienCell, Cat # 1800) were cultured following the manufacturer’s directions using a comprehensive kit (ScienCell). This kit included Astrocyte specialty medium (ScienCell,Cat # 1801), dPBS (ScienCell Cat # 0303), Trypsin EDTA (ScienCell Cat # 0183), Trypsin neutralizing solution, and poly-L-lysine (PLL, ScienCell Cat # 0403). Astrocytes were seeded onto PLL-coated T-75 flasks. The medium was changed every three days until the cultures were approximately 70% confluent. Subsequently, the astrocyte medium was exchanged daily until the culture reached 90% confluence. Astrocytes were then subcultured for experiments and seeded at a density of 5,000 cells/cm².

### Generating 3D forebrain organoids derived from human iPSCs

3D cortical organoid generation followed our prior published methods.^47, 48^ Human iPSC lines and *MAPT* mutation lines (**Table 1**) developed from patient samples and knock-in approaches^95^ were cultured on Matrigel matrix (Corning Cat # 356231) coated 6-well plates (Corning #3506) with mTeSR (StemCell Technologies Cat # 85851). iPSC lines were cultured using FGF2Discs (Stem Cultures Cat # DSC500-48) and fed with mTeSR three times a week. Cultures were expanded and split every 5-7 days when they reached approximately 70-80% confluency using ReLeSR. (StemCell Technologies Cat # 5873). To generate spheroids, iPSCs were first rinsed with DMEM/F12 (Thermo Fisher #11330-032) and then dissociated into a single-cell suspension using Accutase (Thermo Fisher Cat # A1110501), followed by incubation at 37°C in a 95% O_2_/5% CO_2_ incubator for 5 minutes or until the cells detached. The detached cells were transferred to a 15 mL conical tube and counted using a cytoSMART Corning Cell Counter. The cell suspension was then centrifuged at 1,200 rpm for 4 minutes and resuspended in mTeSR medium supplemented with 10 μM ROCK inhibitor Y-27632 (Tocris Cat # 1254) at a concentration of 10,000 cells per 100 µL or 1 million cells per 10 mL. The resuspended cells were transferred to a small trough, and with a multi-channel pipette, gently and evenly distributed at 100 µL per well into a 96-well plate (Prime Surface Cat # MS9096SZ). The plate was then centrifuged at 100 x rcf for 3 minutes. The organoids received daily half-feeds for five days with E6 medium supplemented with freshly prepared aliquots of 2.5 µM Dorsomorphin (DM), 10 µM SB4311541, and 2.5 µM XAV-939 (Tocris Cat # XAV-939). On day 6, the spheroids were transferred to Neuralbasal A medium (ThermoFisher Cat # 10888022) with B27 (without A, #17504044), 1% anti-Anti, 1% GlutaMAX, 20 ng/mL FGF2 (Shenandoah Cat # 100-28), and 20 ng/mL EGF (Shenandoah Cat # 100-26). They were fed daily for 10 days, then every other day for the following 9 days. At 20 days, a subset of organoids was sectioned to confirm proper patterning. This quality control (QC) assessment included markers for dorsal forebrain progenitors such as PAX6, SOX2, FOXG1, and the absence of SOX10. From day 43 onward, the spheroids were fed with the same medium every 4 days, excluding growth factors.

**Table 1.** iPSC lines, mutant and isogenic controls.

| Table 1. iPSC lines, mutant and isogenic controls |  |  |  |
| --- | --- | --- | --- |
|  | Line ID | MAPT genotype | Sex |
| V337M | 1 GIH-6-C1d1E11 | WT/WT (corrected) | F |
|  | 2 GIH-6-C1d1A02 | V337M/WT | F |
|  | 3 GIH-7-C2d2B12 | WT/WT (corrected) | F |
|  | 4 GIH-7-C2d2A01 | V337M/WT | F |
| R406W | 1 F11362.1d1C11 | WT/WT (corrected) | F |
|  | 2 F11362.1d1F10 | R406W/WT | F |
| WT | 1 F11350.1 | WT/WT | M |
| N279K | 1 F11350.1P2B3 | WT/WT | M |
|  | 2 F11350.1P2B5 | N279K/N279K | M |

### Live confocal time-lapse imaging of organoids

Organoids were housed in a live cell imaging humidified chamber with at 5% CO₂, and maintained at 37°C. GFP-Tau-R5L fluorescence in the organoids was imaged using a Zeiss LSM 780 multi- photon confocal microscope. Scans were taken every 12 minutes at a depth of 20-25 µm into the organoids, with each scan lasting 3 minutes. This imaging protocol was carried out continuously over a duration of 3 days.

### Cryostat sectioning, immunostaining, and microscopy

Differentiated 2D cell lines were fixed using 4% paraformaldehyde in PBS for 30 min, then washed three times with 1X PBS. Fixation and cryostat sectioning of 3D organoids was performed as previously described.^47^ Immunostaining was done using antibodies to PAX6 (BioLegend, 1:200), SOX2 (Santa Cruz, 1:100), β-tubulin III (Sigma, 1:1000), FOXG1 (Takara, 1:500), SOX10 (Santa Cruz, 1:100), MAP2AB (Abcam, 1:1000-2000), Total Tau (DAKO Potts, 1:200), DA9 (gift from Peter Davies, 1:1000) or Anti-HA (1:1,000, Cell Signaling). Primary antibodies were incubated overnight at 4°C, washed three times with PBS, and then incubated with corresponding Alexa Fluor conjugated secondary antibody (1:333-1000) for 1 hour at room temperature. Sections were coverslipped and imaged using fluorescence and confocal microscopy (2x Zeiss inverted Axiovert-D1 microscopes with epifluorescence).

### Cell Death assay ethidium homodimer (EthD-1)

A 2mM stock solution of ethidium homodimer-1 (EthD-1) was diluted into 1x phosphate-buffered saline (PBS) to yield a 4 µM working solution. Subsequently, the 4 µM ethD-1/PBS working solution was further diluted into culture medium to achieve a final concentration of 400 nM. The cells were treated with this solution at room temperature for 30 minutes. Medium was aspirated and replaced with 1x PBS solution supplemented with Hoechst dye at a 1:1000 dilution (Molecular Probes Cat # 95355). The cells were incubated with the 1X PBS/Hoechst solution for 5 minutes to counterstain the nuclei. Cells were live-cell immunofluorescence imaged. Cell death was quantified by dividing the number of EthD-positive cells by the total number of nuclei stained with Hoechst, quantified using Cell Profiler™ software.

### Western Blotting

Cell samples were washed with 1X PBS followed by cell lysis using RIPA lysis and extraction buffer (Thermo Fisher Scientific) supplemented with PhosSTOP EASY pack phosphatase inhibitor cocktail tablets and cOmplete protease inhibitor cocktail (Roche Diagostics). Samples were then sonicated for 10 minutes. DC protein assays were performed on samples to generate protein concentration data. From protein assays, sample lysates concentrations were normalized to 1 µg/µL in RIPA lysis buffer plus phosphatase and protease inhibitors and 6X reducing Laemmli SDS-sample buffer (Boston BioProducts, Inc) and heated at 95°C for 5 min to ensure denaturation of proteins. 10 µg of each lysate sample were separated though sodium dodecyl sulfate polyacrylamide gel electrophoresis (SDS-PAGE) using 4-20% Criterion precast gels (Bio-Rad Cat #456-1095). Proteins were blotted onto Polyvinylidene difluoride (PVDF) membranes (Millipore) using a Trans-Blot Turbo Transfer System (Bio-Rad) at 25V for 7min/blot. The PVDF membranes were probed for total tau (DA9; 1:1,000) and GAPDH (as a loading control, Abcam; 1:5,000). HA-tagged intrabodies were probed with monoclonal Anti-HA (1:1,000, Cell Signaling). Samples were normalized to either actin or GAPDH housekeeping proteins with monoclonal antibodies (anti-actin; 1∶1000, Sigma or anti-GAPDH; 1:10,000 Abcam).

ProteinSimple FluorChem E gel/blot digital imager. Densitometry is quantified with Image J software.

### Genome Editing of iPSCs *MAPT* to N279K

Human iPSCs were edited using CRISPR/Cas9 as previously reported.^64^ Briefly, allele specific guide RNA (5’ GCAGATAATTAATAAGAAGCNGG 3’) was designed for the N279K allele, with at least 3bp of mismatch to any other gene in the human genome and validated for activity using the T7E1 assay, which recognizes and cleaves non-perfectly matched DNA. To prepare cells for editing, iPSC colonies were dissociated into single cells via incubation in Accutase for 10 minutes. Single cell iPSC cultures were maintained on Matrigel supplemented with FGF2 stembeads (Stemcultures, SB500) and cultured in mTesR (StemCell Technologies). To edit cells, iPSCs were nucleofected with 1μg pMaxGFP (to assess nucleofection efficiency), 1μg gRNA, 3μg Cas9, and 300μM single stranded oligodeoxynucleotides (ssODN: 5’gggtccagggtggctgtcactcat ccttttttctggctaccaaaggtgcagataattaaGaagaagctggatcttagcaacgtccagccaagtgtggctcaaaggataatatcaaacacgtcc 3’) and the P3 Primary Cell 4D reaction mix (Lonza). We screened a minimum of 96 clones for genetic editing, routinely choosing 2-5 edited clones for further expansion and analysis. Additionally, 1-2 unedited clones that had undergone the genome-editing process but remained unchanged were also selected. Both edited and unedited clones were characterized using qPCR and ICC for pluripotency markers, karyotyping, and Sanger sequencing of on-target and predicted off-target sites. Single-strand DNA sequencing was performed by Azenta Life Sciences to confirm allele-specific sequences.

## Acknowledgements

We are grateful for the encouragement and guidance of our generous colleagues in the tauopathy field, notably in the Tau Consortium and Cure PSP. We thank Drs. Anne Messer and Jeffrey H. Stern, NSCI, for advice throughout the studies. The *MAPT* mutation human iPSC lines were provided through the support of the Tau Consortium of the Rainwater Charitable Foundation. The Neuracell non-profit core facility https://www.neuracell.org/ provided the high-quality iPSCs, neural progenitor cells, and organoids used in the study. We thank the patients and their families and the clinical and research teams for enabling the production of these precious iPSC resources.

We are grateful for the following funding support: NIH: RF1NS123568 (ST, DCB); R35NS097277 (ST), U01AG072464 (ST,TB) R01AG076007 (DCB); AG066444 (CMK).

The Rainwater Charitable Foundation and the Tau Consortium (ST, CMK, DCB). Cure PSP and the South West Florida PSP Support Group (ST, DCB).

AFTD (DCB).

Alzheimer’s Association Research Fellowship, Grant 22-971644 (ESF).

---

**Supplemental Figure 1.**
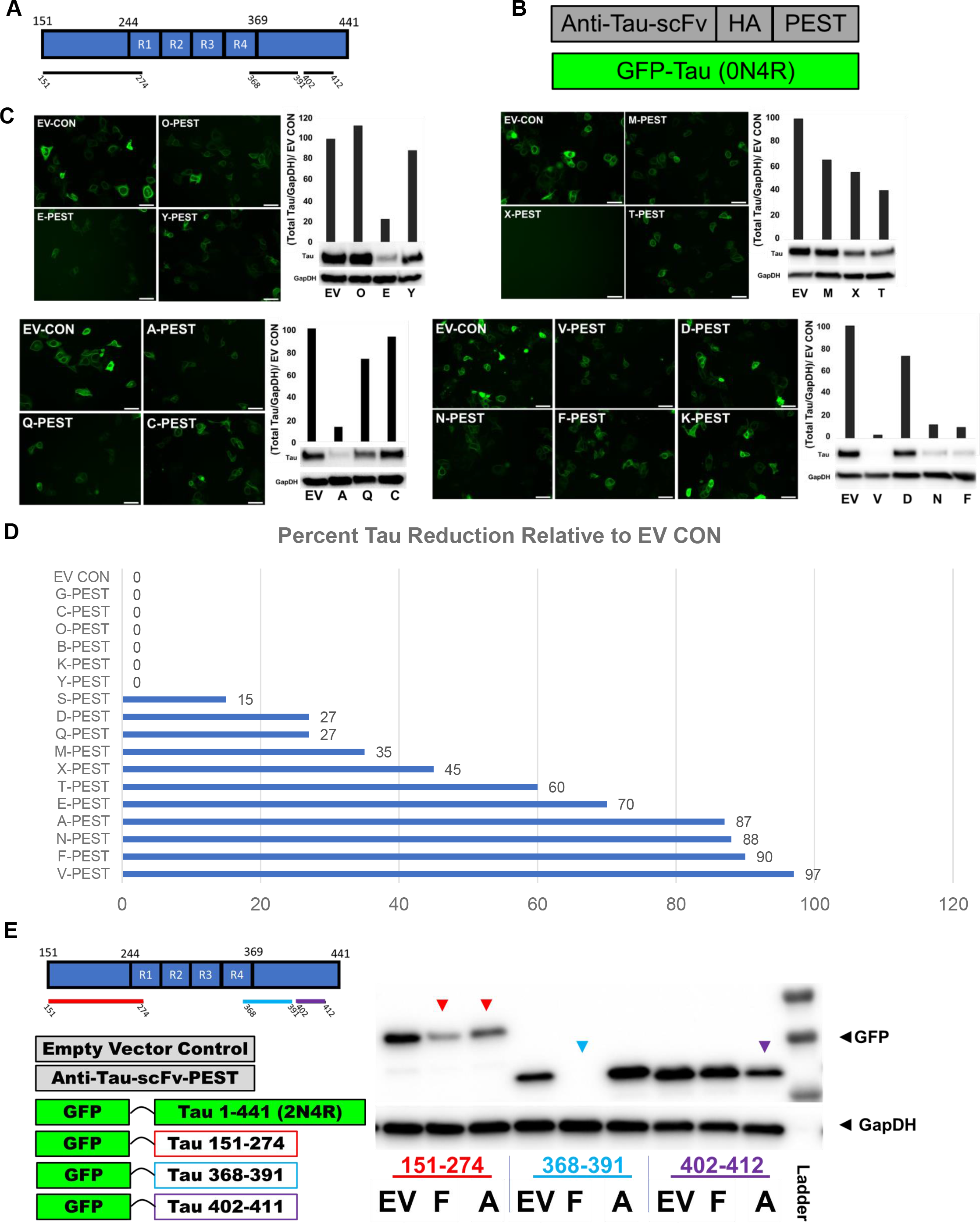
I**d**entification **of bifunctional anti-tau-PEST intrabodies. A)** Antigen binding domain of anti-tau scFvs. **B)** Construct and experimental design. **C)** Representative live cell images. Scale bar = 50 µm and western blot analysis. **D)** Summary of the quantitative western blot analysis. Densitometry bars represent total tau mean optical density values relative to EV CON. **E)** Epitope mapping. Left panel experimental design. Representative western blots show that F-hPEST completely degrades GFP-Tau-368-391 bait compared to EV control. A-PEST and F-PEST appear to partially degrade GFP-Tau-151-274 bait compared to EV control.

**Supplemental Figure 2.**
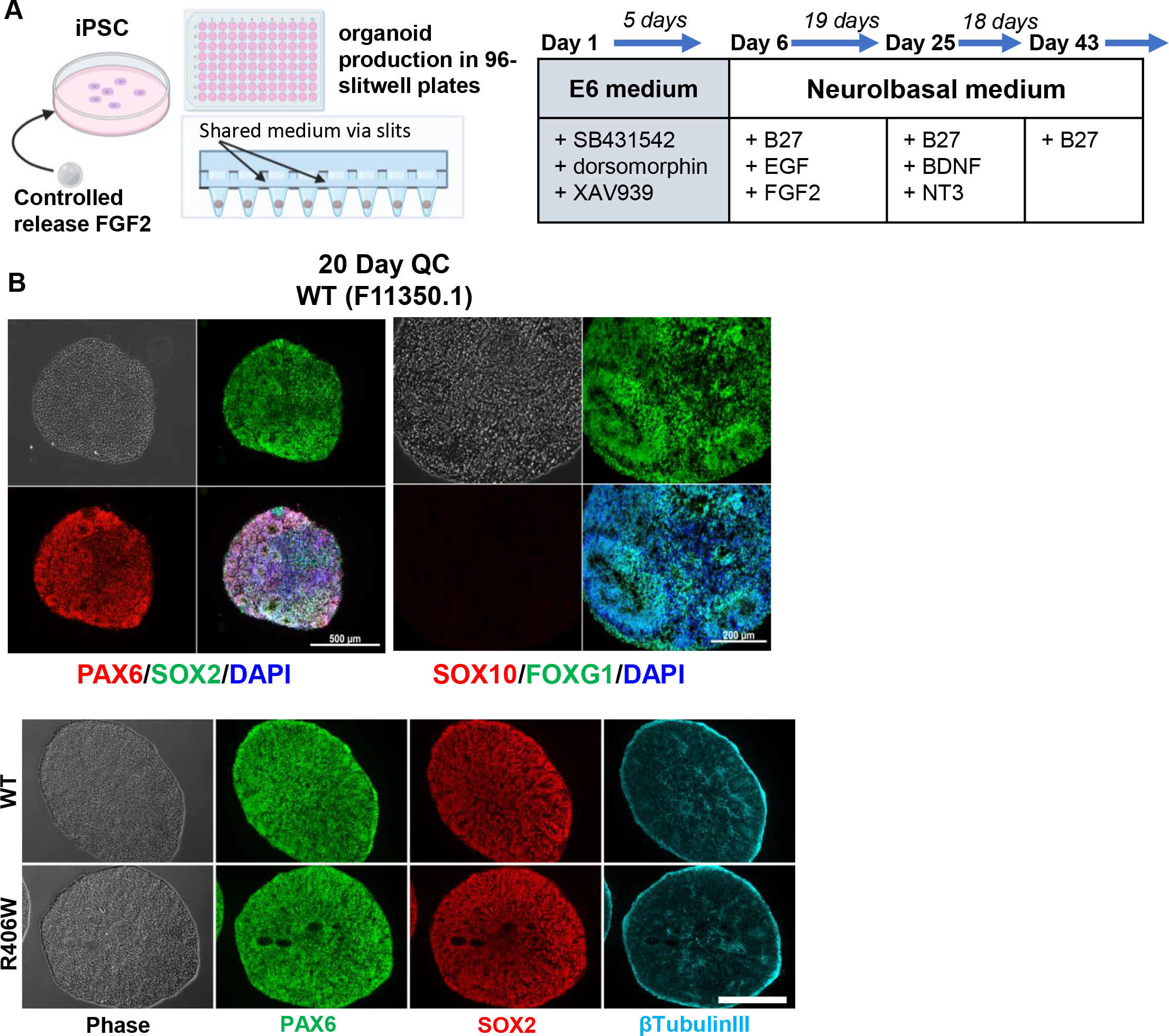
C**o**nfirmation **of cortical organoid patterning. A)** Schematic of iPSC maintenance with stable release FGF2 and 96-slitwell plate method for efficient, scalable, and reproducible cortical organoid production. **B)** Immunofluorescent staining for cortical progenitor markers. At 20 days, organoids express neural stem cell/neuroectoderm markers such as PAX6, SOX2, and FOXG1, and are negative for neural crest marker SOX10. Scale bar = 500 µm.

**Supplemental Figure 3.**
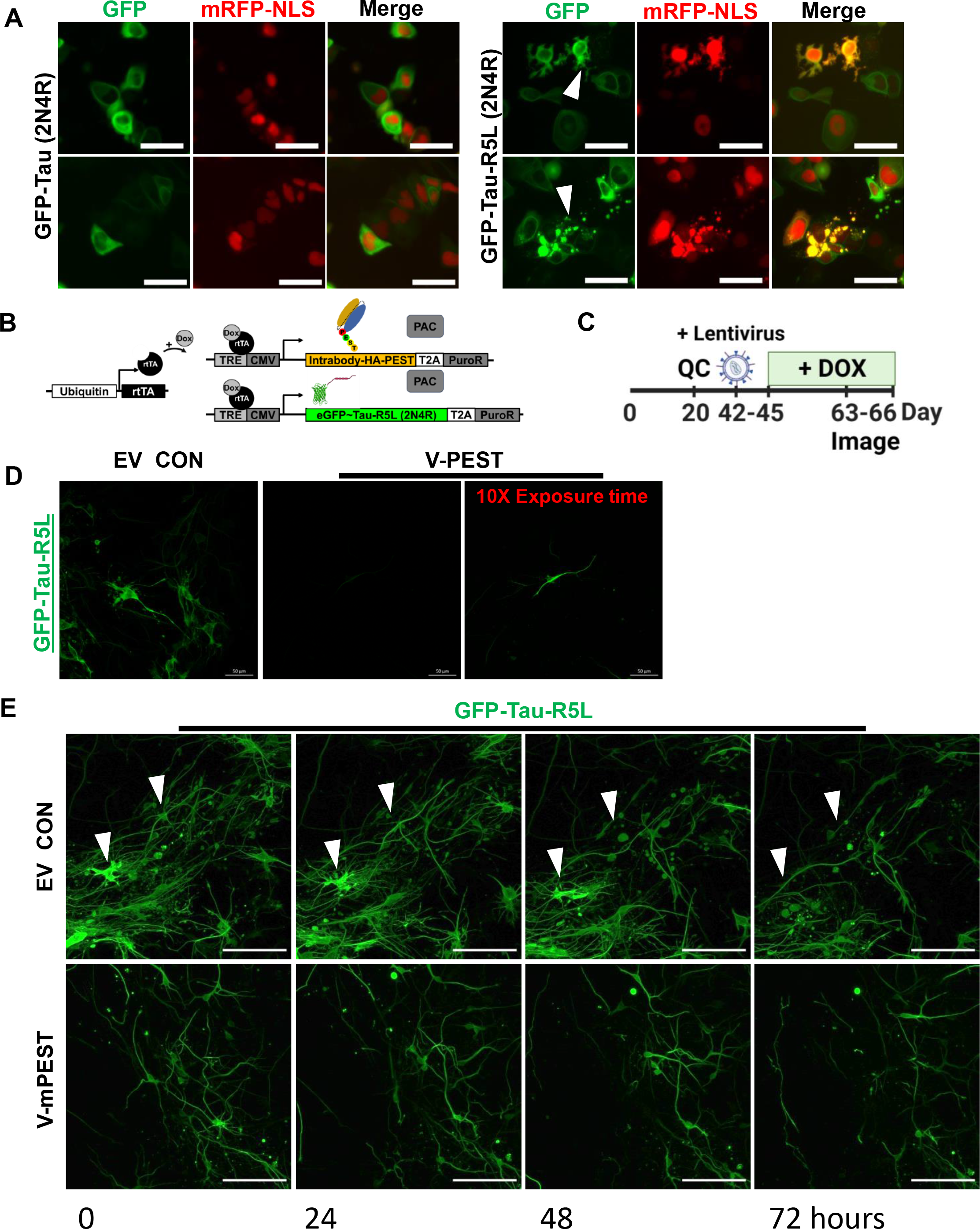
B**i**functional **intrabodies prevent cytotoxicity in 3D organoids that express GFP-Tau-R5L. A)** ST14A murine immortalized cells were co-transfected with either GFP-Tau (2N4R) or GFP-tau-R5L (2N4R) and mRFP-NLS. 72 hours post transfection, cells were live cell imaged for GFP-Tau expression and nuclear mRFP-NLS expression. GFP-Tau-R5L cells contained numerous puncta (arrow; lower panel) and membrane blebbing (arrow; upper panel) Scale bar 50μm**. B)** Schematic of the inducible lentiviral constructs. **C)** Experimental design for intrabody screening in iPSC-derived organoids. **D)** Live confocal imaging of organoids. Following 21 days of treatment with DOX, V-PEST-treated organoids have reduced tau levels and so required a 10x exposure time compared to EV CON to visualize GFP-labeled tau. **E)** GFP-Tau- R5L (2N4R) expression levels were monitored by time-lapse imaging for 3 days. The characteristic morphological signs of apoptosis (Arrows indicate cellular shrinkage and membrane blebbing) can be readily observed in empty control virus-treated organoids compared to anti-tau V-PEST (Scale bar = 50μm).

**Supplemental Figure 4.**
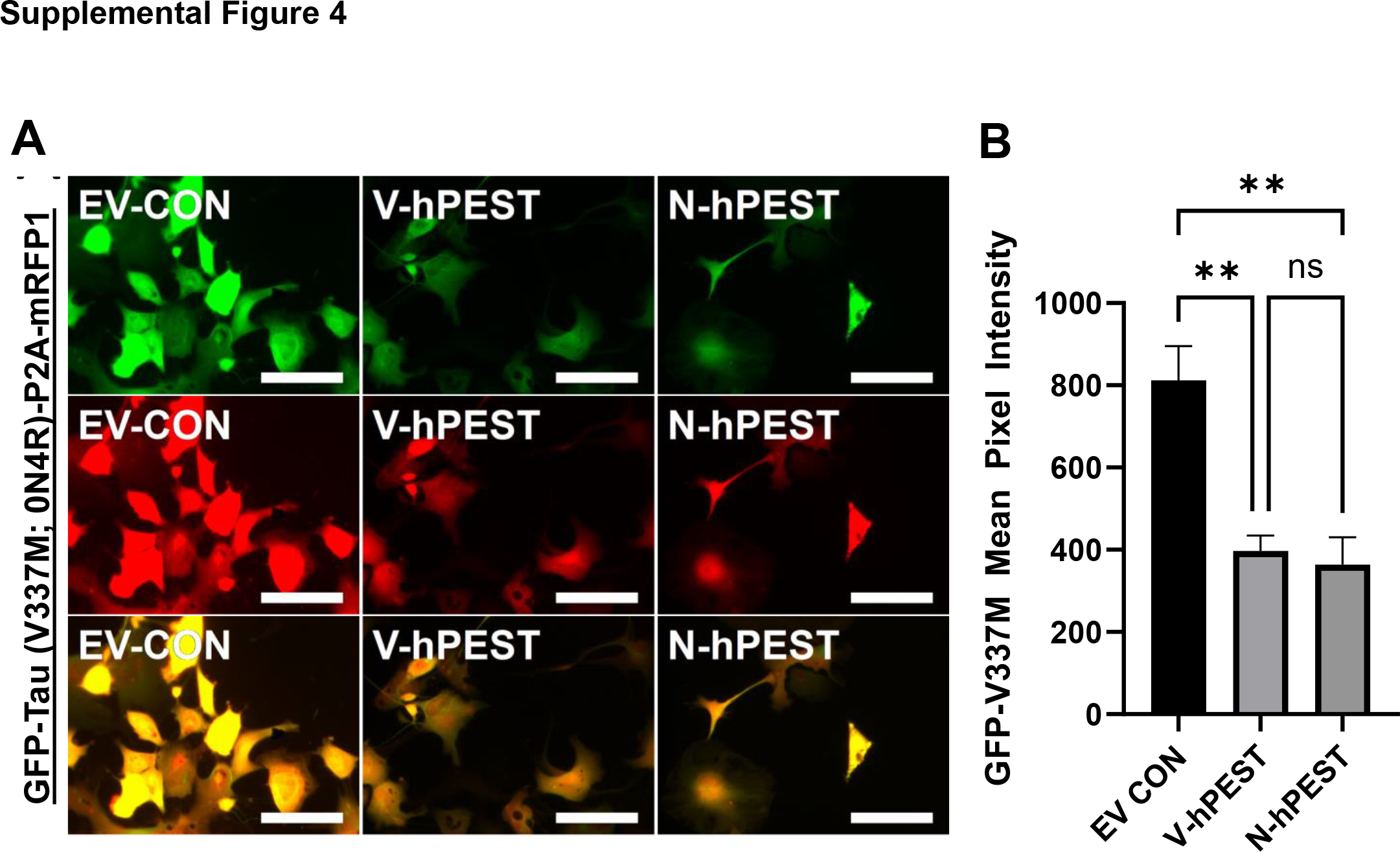
A**n**ti**-tau-hPEST intrabodies V-hPEST and N-hPEST significantly reduced mutant GFP-V337M tau in human astrocytes. A)** Human astrocytes were transduced with GFP-Tau (V337M; 0N4R)-P2A-mRFP1 and either V-hPEST, N-hPEST, or EV CON; live cell images show reduced tau with the V- or N-PEST intrabodies; Scale bar = 100 µm, quantified in **B)** using ImageJ. (Data points, mean+/-SEM, n=3, One-way ANOVA, with Tukey’s post-hoc analysis, ** p < 0.01).

**Supplemental Figure 5.**
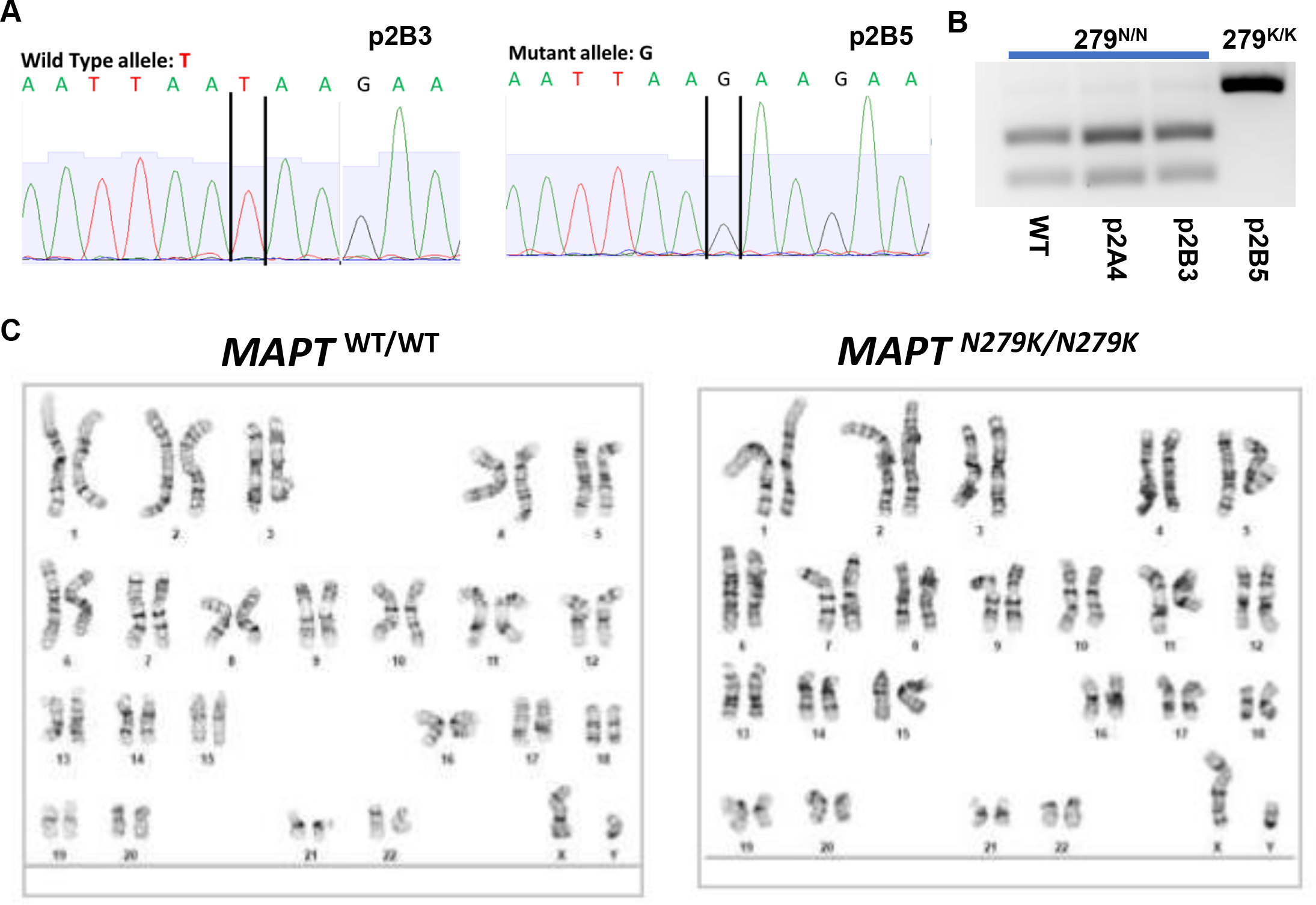
G**e**neration **of N279K isogenic iPSC cell lines.** The N279K mutation was introduced into healthy wild type iPSCs with the following genetic background: Male, Nonwhite hispanic, WT/WT for *MAPT*, APOE E3/E3, H1/H1. **A)** Sanger sequence verification. **B)** Restriction mapping of genomic DNA. Genomic DNA flanking the *MAPT* N279K mutation was digested with Ase1. The Ase1 site is not present in the N279K homozygous DNA. **C).** Karyotyping.

**Supplemental Figure 6.**
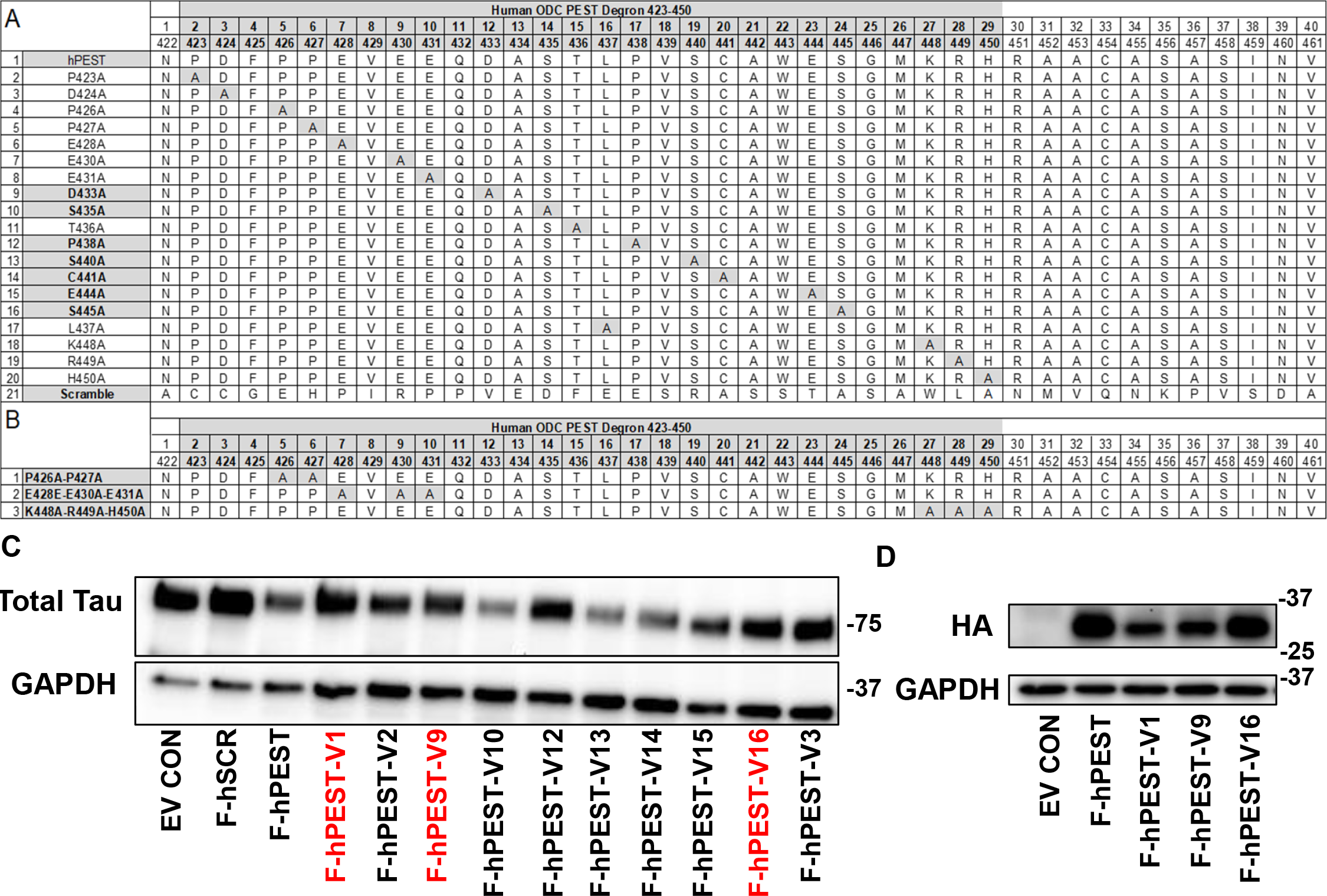
I**d**entification **of anti-tau-PTAP intrabodies. A)** Schematic of human ODC PEST degron and serial alanine mutagenesis of the PEST amino acids. **B)** Compound mutations. Certain single and compound mutations within the hPEST degron (highlighted in grey) are predicted to alter the targeted degradation of the intrabody and its bound antigen. **C-D)** ST14A cells were transfected with either GFP-Tau and either EV-CON or anti-tau PTAP intrabodies. 72 hours post transfection, cells were imaged for GFP fluorescence and then processed for western blotting. **C)** Western blot for total tau (DA9). **D)** Western blotting for intrabody expression (HA).

**Supplemental Video 1. GFP-Tau-R5L treated with EV CON.** Live confocal time-lapse imaging of organoids. Video shows GFP-Tau-R5L cells swelling in processes and undergoing cell death.

**Supplemental Video 2. GFP-Tau-R5L treated with V-mPEST.** Live confocal time-lapse imaging of organoids. Intrabody treated cells appear to be healthy and dynamic without signs of cytotoxicity.

